# Lipid peroxidation intrinsically induces mitochondrial iron overload via Bach1-HO-1 signaling to promote cardiac ferroptosis

**DOI:** 10.1101/2025.10.15.682717

**Authors:** Xiaoyun Guo, Yi Chen, Yachang Zeng, Xiaoliang Mo, Siqi Hong, Hui He, Jing Li, Kevin Zhang, Qinghang Liu

## Abstract

**Background:** Lipid peroxidation and iron accumulation are hallmarks of ferroptosis, a form of cell death characterized by iron-dependent oxidative damage to cellular membranes. However, the molecular link between lipid peroxidation and iron overload in the execution of ferroptosis remains elusive. Moreover, the pathophysiological implications of the interaction between lipid peroxidation and iron overload in cardiac homeostasis and remodeling are also unknown.

**Methods:** We assessed the role of lipid peroxidation in mediating cardiac iron overload and ferroptosis using genetic mouse models. We also performed molecular and cellular biology studies to elucidate the mechanisms by which lipid peroxidation regulates iron homeostasis and ferroptosis signaling in cardiomyocytes.

**Results:** Cardiomyocyte-specific ablation of *Gpx4* (glutathione peroxidase 4), a key suppressor of lipid peroxidation, promoted iron overload and ferroptosis in the heart, leading to dilated cardiomyopathy. Mice with heterozygous *Gpx4* knockout were also predisposed to adverse cardiac remodeling and dysfunction following pressure overload. Mechanistically, elevated lipid peroxidation due to GPX4 inactivation intrinsically induced iron overload by promoting the nuclear export of Bach1 and subsequent induction of heme oxygenase-1 (HO-1). Genetic and pharmacologic inhibition of HO-1 markedly attenuated iron overload and ferroptosis in cardiomyocytes and rescued dilated cardiomyopathy associated with *Gpx4* deficiency. Moreover, we identified HO-1 mitochondrial translocation as a key mechanism driving mitochondrial iron overload and ferroptosis. Targeted inhibition of mitochondrial iron overload or lipid peroxidation abrogated cardiac ferroptosis and pathological remodeling induced by *Gpx4* deficiency.

**Conclusions:** These findings identified a mechanistic link between lipid peroxidation and iron overload via the Bach1-HO-1 signaling pathway, revealing new regulators and molecular targets for cardiac ferroptosis.

## Introduction

Ferroptosis is a form of regulated cell death characterized by iron-dependent accumulation of lipid reactive oxygen species, leading to irreparable lipid damage and membrane permeabilization.^1,2^ Recent studies suggest that ferroptosis plays an important role in the pathogenesis of cardiac pathology induced by pressure overload, ischemic injury, catecholamine overload, doxorubicin cardiomyopathy, and diabetic cardiomyopathy.^3^ Ferroptosis is morphologically and biochemically distinct from other forms of cell death such as apoptosis, necroptosis, and autophagy.^1^ Accumulation of lipid peroxidation products is a hallmark of ferroptosis. Lipid peroxidation is a free radical-mediated process under which reactive oxygen species react with polyunsaturated fatty acids to generate a wide array of oxidation products, such as hydroperoxides and reactive lipid electrophiles. Enhanced lipid peroxidation can be detected in both the cytosol and mitochondria, which plays an important role in the execution of ferroptosis.^2,4–7^ Iron accumulation is another hallmark of ferroptosis, which promotes lipid peroxidation by producing hydroxyl and alkoxyl radicals through the Fenton reaction.^8^ Notably, ferroptotic cell death, regardless of the mechanisms of induction, can be blocked by iron chelators, indicating a central role of iron in the execution of ferroptosis. However, the molecular link between lipid peroxidation and iron accumulation in the execution of ferroptosis remains elusive. Specifically, whether and how lipid peroxidation intrinsically induces iron overload during ferroptosis are poorly understood. Moreover, the pathophysiological implications of the crosstalk between lipid peroxidation and iron overload in cardiac homeostasis and remodeling are also unknown.

Glutathione peroxidase 4 (GPX4) is a key suppressor of lipid peroxidation, which reduces toxic lipid hydroperoxides to the corresponding alcohols by using glutathione as the electron donor, thus preventing oxidative damage to the cells^2^. GPX4 is unique among eight known glutathione peroxides as it is the only enzyme capable of reducing oxidized fatty acids and cholesterol hydroperoxides. Mutations in *GPX4* gene in humans led to premature death by cardiovascular, cerebrovascular, neuromuscular, or renal complications.^9^ Deletion of *Gpx4*, but not other GPX isoforms, caused embryonic lethality in mice, highlighting a non-redundant role for GPX4 (*10,11*). Inducible disruption of *Gpx4* led to acute renal failure and early lethality in mice.^2^ Conditional ablation of GPX4 in neurons resulted in rapid onset of paralysis in the adult mice.^12^ Transgenic overexpression of GPX4 attenuated, whereas heterodeletion of GPX4 exacerbated, doxorubicin-induced cardiomyopathy and myocardial ischemia/reperfusion (I/R) injury.^13,14^ Moreover, GPX4 expression was markedly downregulated in the heart following myocardial infarction, pressure overload, and doxorubicin-induced cardiomyopathy.^13–17^ Although GPX4 as a suppressor of lipid peroxidation is well established, its role in iron homeostasis remains undefined.

Here, we investigated the molecular link between lipid peroxidation and iron accumulation in ferroptosis and its role in cardiac homeostasis and remodeling. We found that cardiomyocyte-specific ablation of *Gpx4,* a key suppressor of lipid peroxidation, triggered iron overload in the heart, leading to dilated cardiomyopathy and premature death. Mechanistically, our data revealed a key link between lipid peroxidation and iron overload via a Bach1-HO-1 dependent mechanism. We also identified HO-1 mitochondrial translocation as a previously undescribed mechanism that drives mitochondrial iron overload and ferroptosis. Importantly, genetic and pharmacologic blockade of Bach1-HO-1 signaling or mitochondrial iron overload rescued cardiac ferroptosis and remodeling, which may serve as new molecular targets for cardiac ferroptosis, pathological remodeling, and heart failure.

## Results

### Enhanced lipid peroxidation via genetic ablation of *Gpx4* induces dilated cardiomyopathy

To investigate the relationship between lipid peroxidation and iron homeostasis and its functional relevance in the heart, we generated a mouse model with cardiomyocyte-specific deletion of *Gpx4*, a key suppressor of lipid peroxidation^2,18^. Mice homozygous for the *Gpx4*-*loxp* targeted allele (*Gpx4^fl/fl^*)^10^ were crossed with αMHC-MerCreMer (MCM) transgenic mice^18^ to ablate GPX4 in cardiomyocytes. Western blot analysis showed that GPX4 was efficiently deleted (> 80%) from the hearts of *Gpx4^fl/fl^*MCM mice but not *Gpx4^fl/fl^* or MCM control mice following tamoxifen treatment (**Fig. 1A**). Strikingly, ∼40% of *Gpx4^fl/fl^*MCM mice showed early lethality within two weeks of tamoxifen administration (**Fig. 1B**). Moreover, histological analysis revealed severe ventricular dilation with high levels of myocardial fibrosis in *Gpx4^fl/fl^*MCM mice, but not *Gpx4^fl/fl^*or MCM controls (**Fig. 1, C** and **D**). The expression of cardiac fetal genes, including *Nppa*, *Nppb*, and *Acta1*, was also greatly elevated in *Gpx4*-deficient hearts (**Fig. 1E**). Echocardiographic analysis showed severe ventricular dilation and contractile dysfunction in *Gpx4^fl/fl^*MCM mice, as indicated by decreased fractional shortening (FS) and increased left ventricular dimensions (LVED and LEVS) (**Fig. 1, F** to **I**). *Gpx4^fl/fl^*MCM mice also displayed enhanced cardiac hypertrophy compared with littermate controls as assessed by increased heart weight to body weight (HW/BW) ratio (**Fig. 1J**). These results indicate that loss of GPX4 promotes adverse cardiac remodeling and dysfunction, suggesting a key homeostatic role for GPX4 in the adult myocardium.

**Figure 1.**
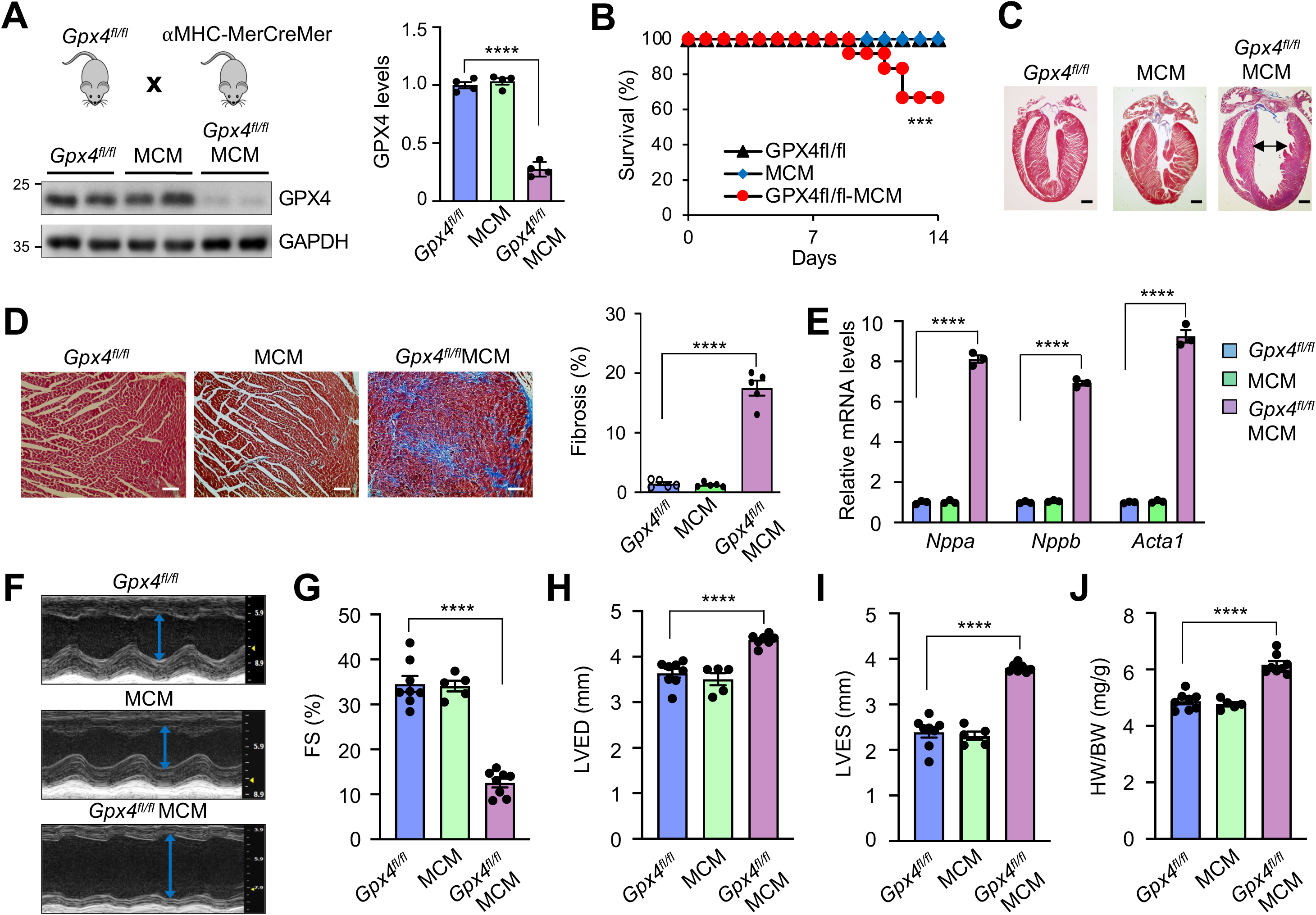
Enhanced lipid peroxidation with genetic ablation of *Gpx4* induces dilated cardiomyopathy and heart failure. (**A**) Western blotting for GPX4 in ventricular extracts from *Gpx4^fl/fl^*, MCM, and *Gpx4^fl/fl^*-MCM mice 2 weeks after tamoxifen administration as described in Methods. GPX4 protein levels were normalized for GAPDH. n = 4. (**B**) Survival rate of the indicated mice treated as in A. n = 5-8. (**C** and **D**) Masson’s trichrome-stained cardiac sections from mice as described in A. Scale bars, 1 mm in C and 50 μm in D. Myocardial fibrosis was quantified with MetaMorph software. n = 5-7. (**E**) mRNA levels of atrial natriuretic peptide (*Nppa*), b-type natriuretic peptide (*Nppb*), and skeletal muscle α-actin (*Acta1*) from cardiac extracts of the indicated genotypes. n = 3. (**F**) Representative echocardiographic M-mode images from mice as described in A. The arrowed lines indicate left ventricular dimension in end-diastole (LVED). (**G-I**) Echocardiographic assessment of fractional shortening (FS) and left ventricular dimension in end-diastole (LVED) and end-systole (LVES) from mice of the indicated genotypes. n = 5-8. (**J)** Heart weight to body weight ratio (HW/BW) from the indicated genotypes. n = 5-8. *\*\*\*P* ≤ 0.001; *\*\*\*\*P* ≤ 0.0001. Statistical analysis was performed using one-way ANOVA with Tukey’s post hoc test (A, D, E, G-J) or log-rank test (B).

It has been shown that GPX4 expression is markedly downregulated in the heart following myocardial infarction, I/R injury, and doxorubicin-induced cardiomyopathy.^13–17^ This prompted us to further examine the role of GPX4 in cardiac remodeling and heart failure propensity in response to pathological stimulation. Given that ablation of GPX4 in *Gpx4*^fl/fl^MCM mice induced severe cardiac remodeling and failure, mice heterozygous for *Gpx4*-loxP allele with αMHC-MerCreMer (*Gpx4*^fl/+^MCM) were used to evaluate the role of GPX4 in pressure overload-induced cardiomyopathy. In contrast to the *Gpx4*^fl/fl^MCM mice, *Gpx4*^fl/+^MCM mice were overtly normal, with no signs of cardiac remodeling or dysfunction at baseline as assessed by echocardiography and histological analysis (**Fig. S1**). However, *Gpx4*^fl/+^MCM mice displayed exacerbated cardiac remodeling and ventricular dilation after 2 weeks of transverse aortic constriction (TAC), with a greater loss in cardiac contractile performance, greater cardiac fibrosis, and more prominent cardiac hypertrophy and lung congestion compared with MCM controls (**Fig. S1**). Therefore, mice with *Gpx4* deficiency were predisposed to adverse cardiac remodeling and heart failure following pressure overload, suggesting a critical cardioprotective role for GPX4 in response to pathological stress.

### Loss of GPX4 promotes cardiac iron overload and ferroptotic cell death via an HO-1 dependent mechanism

We next assessed whether ablation of *Gpx4* induces iron overload and ferroptotic cell death in the heart. Indeed, non-heme iron levels were markedly elevated in the heart of *Gpx4*^fl/fl^MCM (KO) mice compared with αMHC-MerCreMer (MCM) controls, with no significant changes in total cardiac iron levels (**Fig. 2A**). Conversely, the heme levels were significantly decreased in *Gpx4*-deficient mice (**Fig. 2A**). Strikingly, Western blot analysis revealed that heme oxygenase-1 (HO-1), a key enzyme that catalyzes iron release via heme degradation, was greatly elevated in the *Gpx4*-deficient hearts (**Fig. 2B**). Of note, the expression of NAD(P)H quinone oxidoreductase 1 (NQO1), another NRF2-regulated antioxidant enzyme, was unaffected (**Fig. 2B**). These observations led us to hypothesize that *Gpx4* deficiency promotes cardiac iron accumulation and ferroptosis through an HO-1 dependent mechanism. To test this hypothesis, we first determined whether inactivation of HO-1 could rescue the pathological cardiac phenotype associated with *Gpx4*-deficiency. *Gpx4*^fl/fl^MCM (KO) and MCM control mice were administrated with zinc protoporphyrin IX (ZnPP), an HO-1 inhibitor, along with tamoxifen for 2 weeks (**Fig. 2C**). Evans blue dye (EBD) uptake, an indicator of membrane rupture and cell death, was markedly increased in the *Gpx4*-deficient hearts, which was effectively ameliorated by ZnPP treatment (**Fig. 2D**). A similar effect was obtained with Ferrostatin-1 (Fer-1), a classic ferroptosis inhibitor, indicating ferroptosis was induced in the *Gpx4*-deficienct heart in vivo (**Fig. 2D**). Moreover, the serum levels of cardiac troponin I (cTnI), a biomarker for myocardial injury, were also elevated in *Gpx4*-deficient mice, which was also diminished by Fer-1 or ZnPP (**Fig. 2E**). Intriguingly, the number of TUNEL positive cells was greatly increased in the *Gpx4*-deficient hearts, which was also inhibited by Fer-1 or ZnPP (**Fig. 2F**). GPX4 silencing was sufficient to induce ferroptotic cell death in neonatal cardiomyocytes (**Fig. S2**). Together, these data suggest that loss of GPX4 promotes iron accumulation and ferroptotic cell death in the myocardium through an HO-1 dependent mechanism.

**Figure 2.**
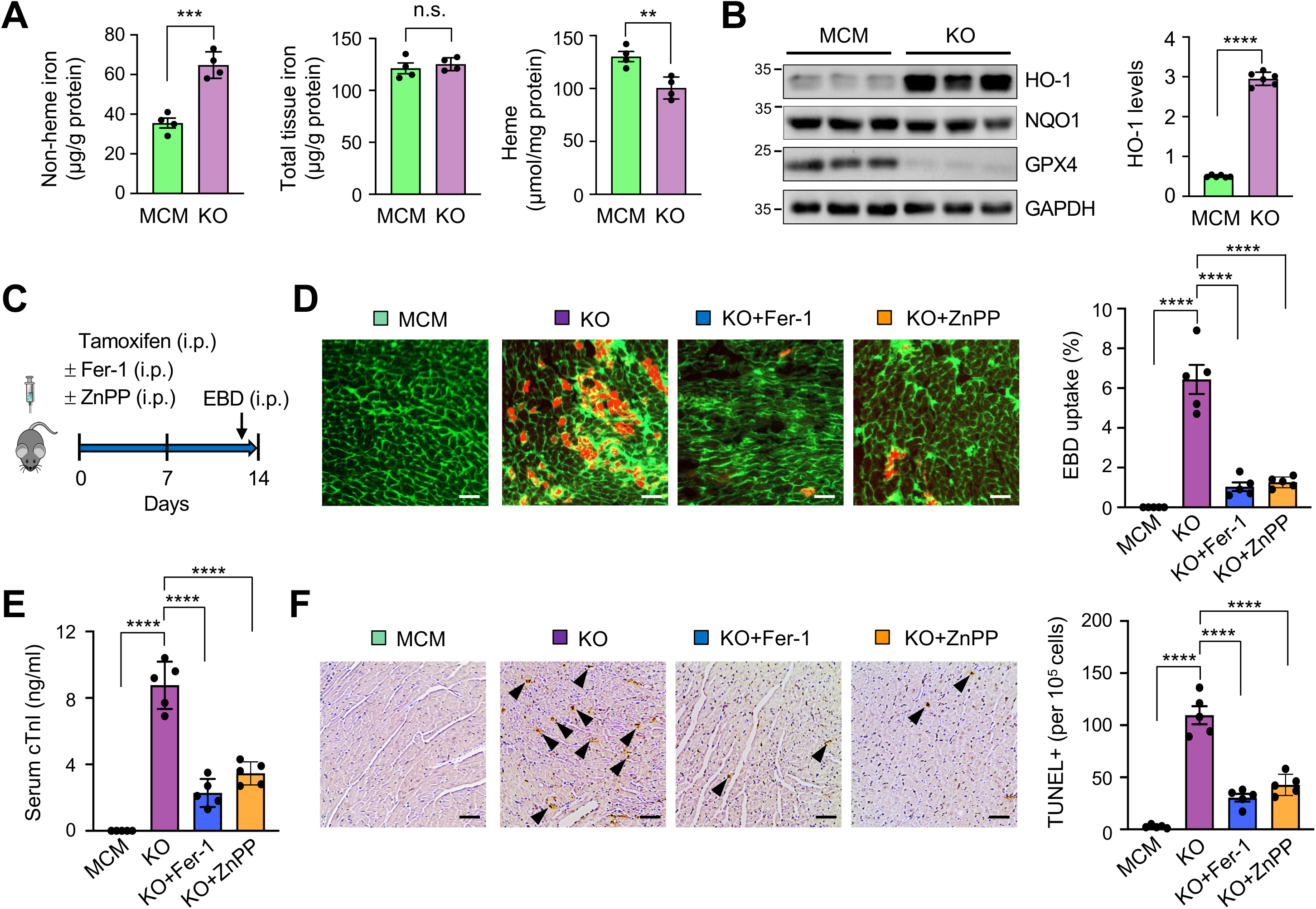
Loss of GPX4 promotes cardiac iron overload and ferroptotic cell death via an HO-1 dependent mechanism. (**A**) Non-heme iron, total iron, and heme levels from cardiac extract of MCM and *Gpx4^fl/fl^*-MCM (KO) mice. n = 4. (**B**) Western blot and quantification of HO-1 from cardiac extract of MCM and KO mice. n = 6. (**C**) Experimental regimen for the administration of tamoxifen along with Fer-1 (1 mg/kg/day, i.p.) or ZnPP (5 mg/kg/day, i.p.) followed by EBD injection 24 h before euthanasia. (**D**) Representative cardiac sections imaged for wheat germ agglutinin (WGA; green fluorescence) and EBD uptake into cardiomyocytes (red fluorescence) from mice subjected to the experimental regimen shown in A. Scale bar, 50 μm. Quantified data indicate EBD positive areas from the indicated groups. n = 5. (**E**) Serum cardiac troponin I (cTnI) levels from the indicated groups. n = 5. (**F**) TUNEL staining and quantification in cardiac sections from the indicated groups. Arrow heads indicate TUNEL positive cells. Scale bar, 50 μm. n = 5. *\*\*P ≤* 0.01; *\*\*\*P ≤* 0.001; *\*\*\*\*P ≤* 0.0001; n.s., not significant. Statistical analysis was performed using Student’s t-test (A, B) or one-way ANOVA with Tukey’s post hoc test (D, E, F).

### Inactivation of HO-1 attenuates cardiac ferroptosis and remodeling associated with *Gpx4*-deficiency

We next examined whether blockade of lipid peroxidation or inactivation of HO-1 rescues cardiac remodeling and dysfunction in *Gpx4* knockout (KO) mice. Strikingly, treatment with Fer-1 and ZnPP both prevented early lethality of *Gpx4*^fl/fl^MCM (KO) mice following tamoxifen administration (**Fig. 3A**). Fer-1 largely rescued cardiac dysfunction, ventricular dilation, cardiac fibrosis, and hypertrophy in *Gpx4*-deficient mice (**Fig. 3, B** to **H**). Moreover, the levels of *ptgs2* mRNA, a signature gene of ferroptosis, and malondialdehyde (MDA), a marker of lipid peroxidation, were greatly increased in *Gpx4*-deficient hearts, which were also abrogated by Fer-1 (**Fig. 3, I** and **J**). These results suggest that the pathological cardiac phenotype associated with *Gpx4* deficiency is primarily driven by ferroptosis. Moreover, cardiac pathology in *Gpx4*-deficient mice was also effectively alleviated by HO-1 inactivation with ZnPP (**Fig. 3, B to J**), revealing a key pathogenic role of HO-1 in this process.

**Figure 3.**
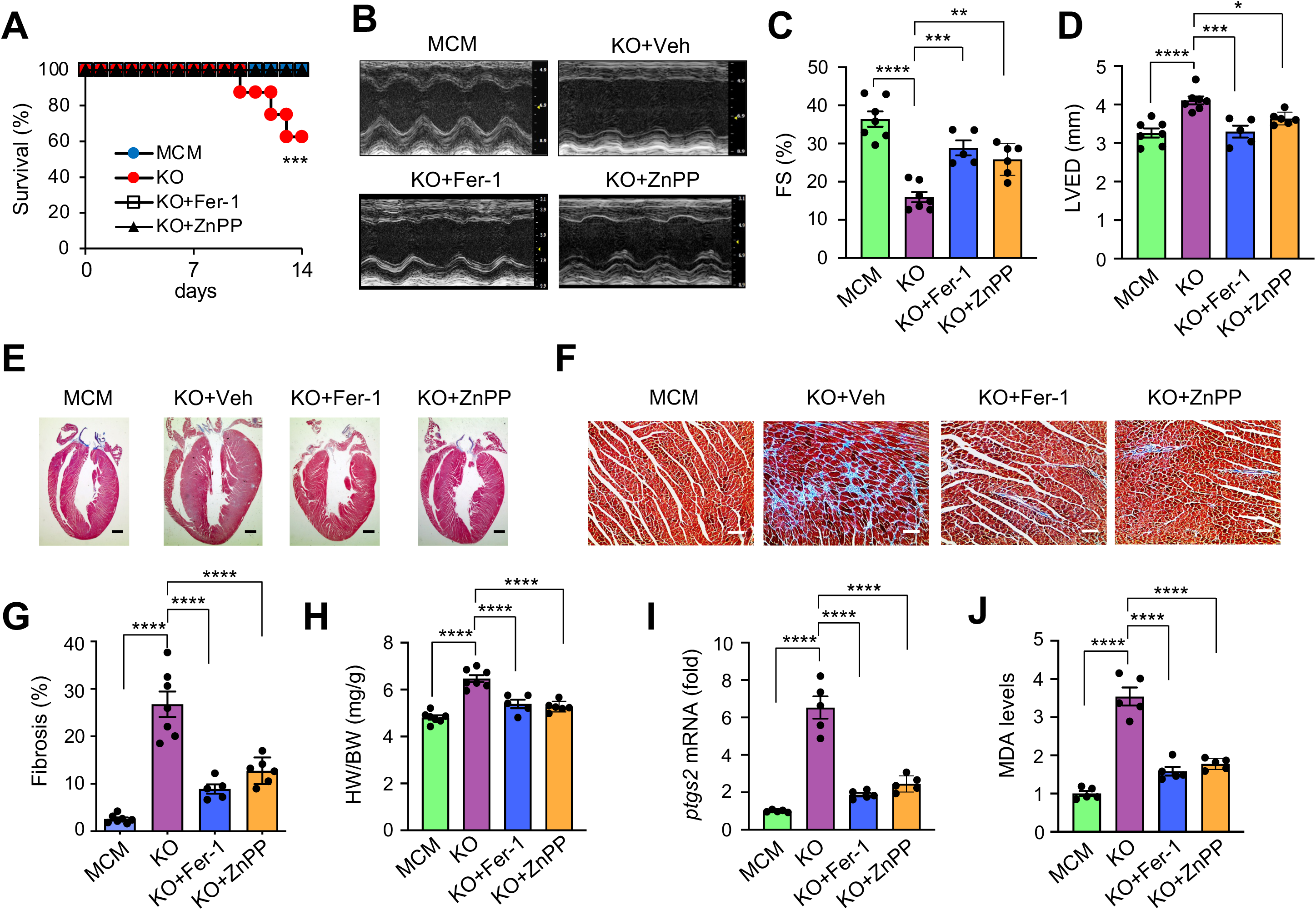
Blockade of lipid peroxidation or inhibition of HO-1 attenuates cardiac remodeling and dysfunction in *Gpx4* deficient mice. (**A**) Survival rate of the indicated mice administrated with tamoxifen along with Fer-1 (1 mg/kg/day, i.p.) or ZnPP (5 mg/kg/day, i.p.) for 2 weeks. (**B**) Representative M-mode echocardiographic images from mice treated as in A. (**C** and **D**) Fractional shortening (FS) and left ventricular end-diastolic dimension (LVED) measured by echocardiography from mice treated as in A. n = 5-7. (**E** and **F**) Low-and high-magnification images of Masson’s trichrome-stained cardiac sections from mice treated as in A. Scale bars: 1 mm in E; 50 μm in F. (**G**) Cardiac fibrosis quantified with MetaMorph software. n = 5-7. (**H**) Heart weight to body weight ratio (HW/BW) from the indicated mice. n = 5-7. (**I**) *ptgs2* mRNA levels in the heart from the indicated groups. n = 5-7. (**J**) MDA levels from cardiac extracts from the indicated groups. n = 5-7. *\*P ≤* 0.05; *\*\*P ≤* 0.01; *\*\*\*P ≤* 0.001; *\*\*\*\*P ≤* 0.0001. Statistical analysis was performed using log-rank test (A) or one-way ANOVA with Tukey’s post hoc test (C, D, G-J).

### Bach1-HO-1 signaling regulates cardiomyocyte ferroptosis

We further examined whether GPX4 regulates ferroptosis via HO-1 in cardiomyocytes in vitro. Consistent with our in vivo finding above, HO-1 expression in cardiomyocytes was also upregulated by GPX4 silencing (**Fig. 4A**). In contrast, the expression of NQO1, another antioxidant protein, was unaffected (**Fig. 4A**). Importantly, labile iron levels in cardiomyocytes were markedly increased by GPX4 deletion, which was reversed by HO-1 deletion (**Fig. 4B**). Consistent with this, cell death induced by GPX4 deletion was also inhibited by HO-1 deletion (**Fig. 4B**). Moreover, inhibition of HO-1 with ZnPP effectively inhibited cardiomyocyte ferroptosis induced by RSL3, a specific GPX4 inhibitor^18^ (**Fig. S3**). Conversely, hemin, an HO-1 activator, exaggerated RSL3-induced ferroptosis (**Fig. S3**). We further examined whether the RSL3-induced ferroptosis is regulated by HO-1 levels, given the controversial effects of HO-1 in previous studies.^20^ Adenoviral vector mediated HO-1 knockdown clearly elicited an anti-ferroptotic effect in cardiomyocytes. However, this effect was gradually lost with further HO-1 depletion (**Fig. 4C**). Therefore, our results revealed a biphasic effect of HO-1 on RSL3-induced ferroptosis in cardiomyocytes.

**Figure 4.**
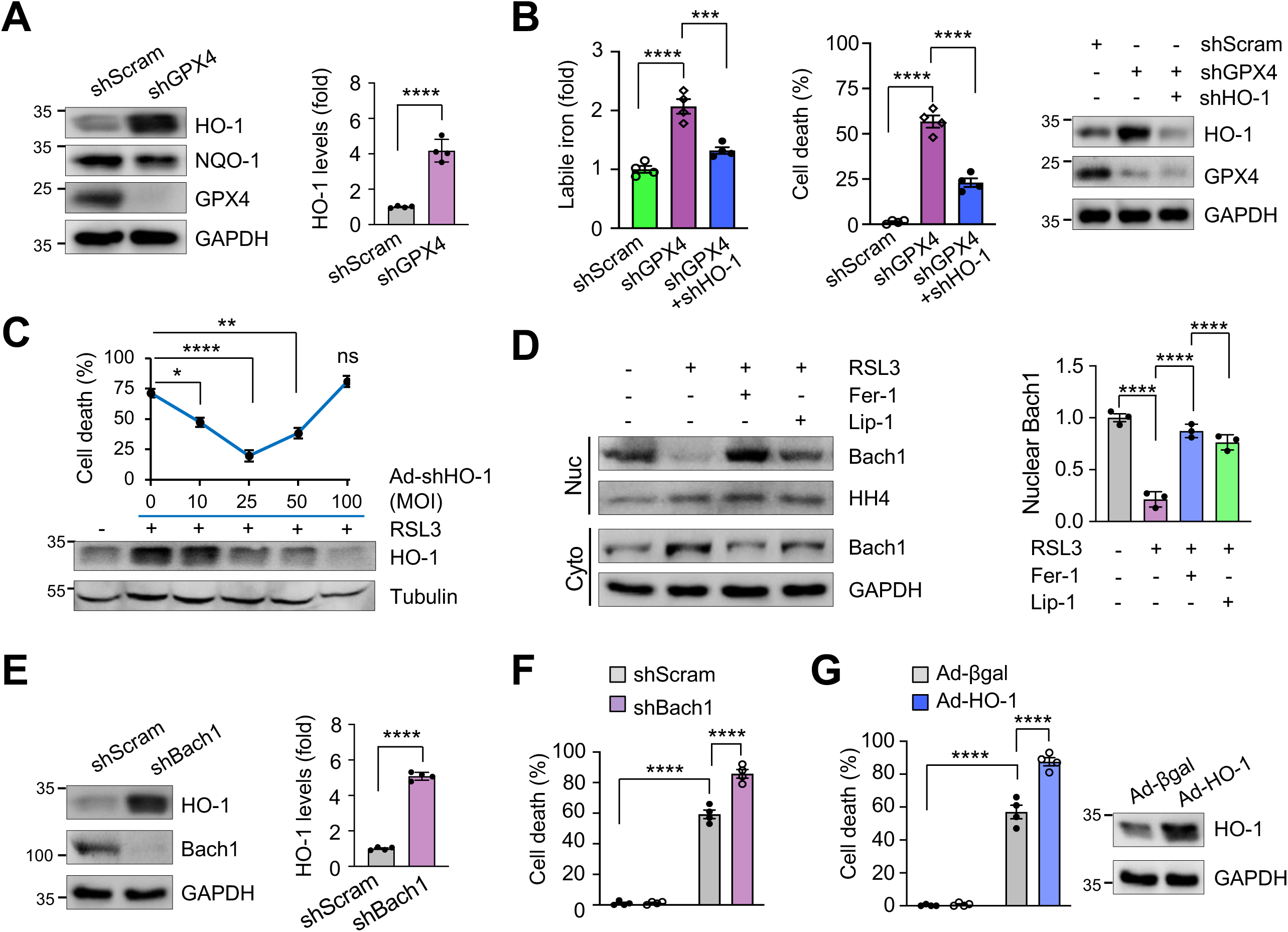
Inhibition of GPX4 induces iron accumulation and ferroptosis via a Bach1-HO-1 dependent mechanism. (**A**) Western blotting for the indicated proteins in neonatal cardiomyocytes infected with adenoviral vectors encoding GPX4 shRNA (shGPX4) or a scrambled sequence (shScram) for 48 h. n = 4. (**B**) Labile iron levels and cell death in neonatal cardiomyocytes treated as indicated for 72 h. shHO-1, adenoviral vector encoding HO-1 shRNA. n = 4. (**C**) Cell death in neonatal cardiomyocytes infected with Ad-shHO-1 at the indicated MOIs followed by 0.5 μM RSL3 for 12 h. n = 4. (**D**) Western blotting for the indicated proteins from nuclear and cytosolic fractions of neonatal rat cardiomyocytes treated with 0.5 μM RSL3 in the presence of 10 μM Fer-1 or 1 μM Lip-1 for 12 h. n = 3. (**E**) Western blotting and quantification of HO-1 and Bach1 in neonatal rat cardiomyocytes infected with Ad-shBach1 or Ad-shScram for 24 h. n = 4. (**F**) Cell death in neonatal rat cardiomyocytes infected with the indicated vectors followed by treatment with 0.5 μM RSL3 or vehicle control for 12 h. n = 4. (**G**) Cell death in neonatal rat cardiomyocytes infected with Ad-βgal or Ad-HO-1 for 24 h followed by treatment with 0.5 μM RSL3 or vehicle control for 12 h. n = 4. *\*P ≤* 0.05; *\*\*P ≤* 0.01; *\*\*\*P ≤* 0.001; *\*\*\*\*P ≤* 0.0001. Statistical analysis was performed using Student’s t-test (A, E), one-way ANOVA with Tukey’s post hoc test (B, C, D), or 2-way ANOVA with Tukey’s post hoc test (F, G).

To determine the mechanism by which GPX4 regulates HO-1 signaling, we found that Bach1, a transcriptional suppressor of HO-1 (*21*), was translocated from the nucleus to the cytosol upon RSL3 stimulation (**Fig. 4D**). This was associated with a decease in total cellular levels of Bach1 (**Fig. 3C**), likely mediated by proteasomal degradation.^21^ Intriguingly, blockade of lipid peroxidation with Fer-1 or Lip-1 largely abrogated RSL3-induced Bach1 translocation, supporting the notion that Bach1 is regulated by redox states^22^ (**Fig. 4D**). Moreover, RSL3 mediated HO-1 upregulation was also blocked by Fer-1 or Lip-1 (**Fig. S3**). We further determined that deletion of Bach1 was sufficient to enhance HO-1 expression and RSL3-induced ferroptosis in cardiomyocytes (**Fig. 4, E** and **F**). Conversely, overexpression of Bach1 inhibited RSL3-induced ferroptosis (**Fig. S3**). In line with this, overexpression of HO-1 led to exaggerated ferroptosis in cardiomyocyte (**Fig. 4G**). In addition to Bach1, HO-1 is also potentially regulated by the transcription factor NRF2. However, in contrast to the pro-ferroptotic effect of HO-1 described above, overexpression of NRF2 inhibited RSL3-induced ferroptosis in cardiomyocytes, whereas deletion of NRF2 enhanced it (**Fig. S4**). These observations suggest that GPX4 regulates HO-1 expression and ferroptotic cell death mainly through a Bach1-dependent mechanism. Together, these results identify a key role of Bach1-HO-1 signaling in mediating ferroptosis induced by GPX4 inactivation. Moreover, we also examined the role of NCOA4-mediated ferritinophagy in RSL3-induced ferroptosis in cardiomyocytes. Deletion of NCOA4 had minimal effects, suggesting a dispensable role of NCOA4-mediated ferritinophagy in this process (**Fig. S5**).

### Genetic ablation of *Hmox1* inhibits cardiomyocyte ferroptosis, cardiac remodeling and dysfunction induced by *Gpx4* deficiency

To validate the role of HO-1 in mediating cardiac ferroptosis and pathological remodeling in *Gpx4*-deficient mice in vivo, HO-1 (encoded by *Hmox1* gene) was genetically ablated by crossing *Hmox1*-floxed mice^23^ with *Gpx4*^fl/fl^MCM mice. To circumvent the potential detrimental effects of HO-1 ablation in homozygous *Hmox1*^fl/fl^MCM mice, here we used heterozygous *Hmox1*^fl/+^MCM to partially inactivate HO-1 in *Gpx4*-deficient mice. Mice of different genotypes were confirmed by Western blot analysis of HO-1 and GPX4 protein expression in the heart (**Fig. 5A**). Consistent with our results using the HO-1 inhibitor ZnPP, genetic deletion of HO-1 also largely rescued the pathological phenotype of *Gpx4*-deficient mice by preventing cardiac fibrosis, contractile dysfunction, ventricular dilation, and cardiac hypertrophy (**Fig. 5, B** to **F**). Moreover, HO-1 ablation also attenuated cardiac *ptgs2* mRNA and MDA levels in *Gpx4*-deficient mice (**Fig. 5, G** and **H**). These data further confirm a pathogenic role of HO-1 in mediating adverse cardiac remodeling and dysfunction triggered by *Gpx4* deficiency.

**Figure 5.**
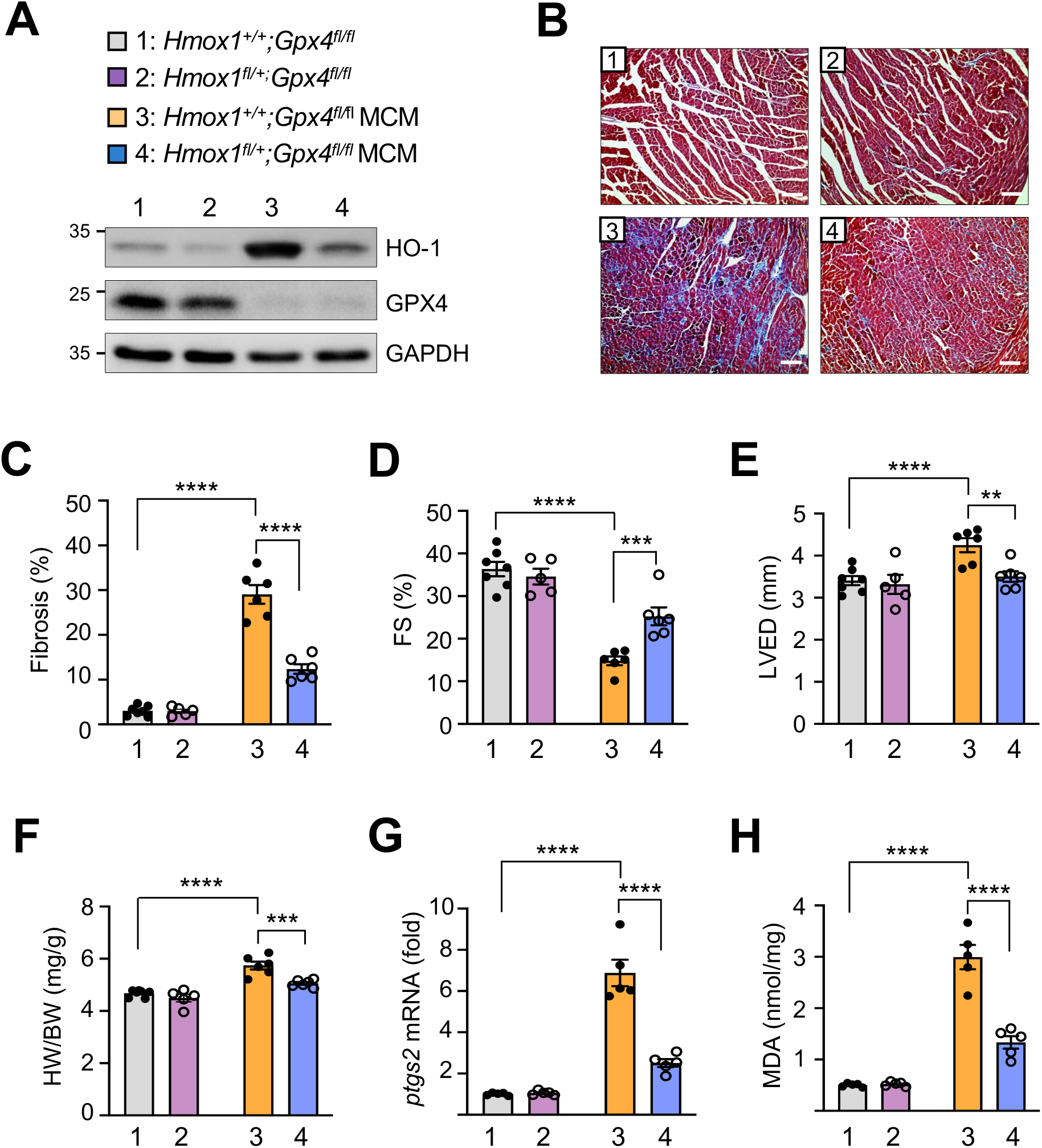
Genetic ablation of *Hmox1* prevents cardiac ferroptosis and remodeling induced by *Gpx4* deficiency. (**A**) Western blotting for HO-1 and GPX4 in cardiac extracts from mice of the indicated genotypes 2 weeks after tamoxifen administration. (**B**) Masson’s trichrome-stained cardiac sections from mice of the indicate genotypes. Scale bar, 50 μm. (**C**) Myocardial fibrosis was quantified by MetaMorph software. n = 5-6. (**D** and **E**) FS and LVED measured by echocardiography from mice of the indicated genotypes 2 weeks after tamoxifen administration. n = 5-6. (**F**) Heart weight to body weight ratio (HW/BW) from the indicated genotypes. n = 5-6. (**G**) *ptgs2* mRNA levels in the heart of the indicated groups. n = 5. (**H**) MDA levels in cardiac extracts from the indicated groups. n = 5. *\*\*P ≤* 0.01; *\*\*\*P ≤* 0.001; *\*\*\*\*P ≤* 0.0001. Statistical analysis was performed using 2-way ANOVA with Tukey’s post hoc test (C-H).

### HO-1 translocation from the cytosol to mitochondria mediates mitochondrial iron accumulation

Intriguingly, we found that mitochondrial ferrous iron (Fe^2+^) levels in cardiomyocytes were markedly increased upon GPX4 inactivation with RSL3, as assessed using mito-FerroGreen, a mitochondria-targeted Fe^2+^ probe (**Fig. 6A**). A significant translocation of HO-1 from the cytosol to mitochondria was also detected in cardiomyocytes following RSL3 treatment (**Fig. 6B**). Moreover, RSL3-induced mitochondrial Fe^2+^ accumulation was largely abrogated by HO-1 deletion, suggesting a key role for HO-1 in mediating mitochondrial iron overload (**Fig. 6C**). To further determine the relationship between mitochondrial iron overload and ferroptosis, cardiomyocytes were infected with an adenoviral vector encoding mitochondrial ferritin (FTMT), a mitochondrial matrix protein that chelates and oxidizes iron. Indeed, RSL3-induced mitochondrial iron accumulation was greatly inhibited by FTMT (**Fig. 6D**). Moreover, RSL3-induced mitochondrial lipid peroxidation was also alleviated, as assessed using mitoPeDPP (**Fig. 6E**). Importantly, overexpressing FTMT largely blocked RSL3-induced cell death in cardiomyocytes (**Fig. 6F**), whereas deletion of FTMT aggravated it (**Fig. S6**), suggesting that mitochondrial iron overload plays a key role in ferroptosis induced by GPX4 inactivation. Moreover, the levels of mitochondrial ROS, as measured by mitoSOX, were also greatly elevated upon RSL3 treatment (**Fig. 6G**). Overexpression of the mitochondria-targeted catalase (mCAT) largely inhibited RSL3-induced mitochondrial ROS accumulation and cell death (**Fig. 6, G** and **H**). Moreover, mitochondria-targeted ROS scavengers^24^, including mitoquinone mesylate (mitoQ) and SKQ1, also inhibited RSL3-induced mitochondrial ROS accumulation, lipid peroxidation, and cell death in cardiomyocytes (**Fig. 6, I** to **K**). Together, these results identified mitochondrial iron overload and lipid peroxidation as key mediators of ferroptosis in cardiomyocytes induced by GPX4 inactivation.

**Figure 6.**
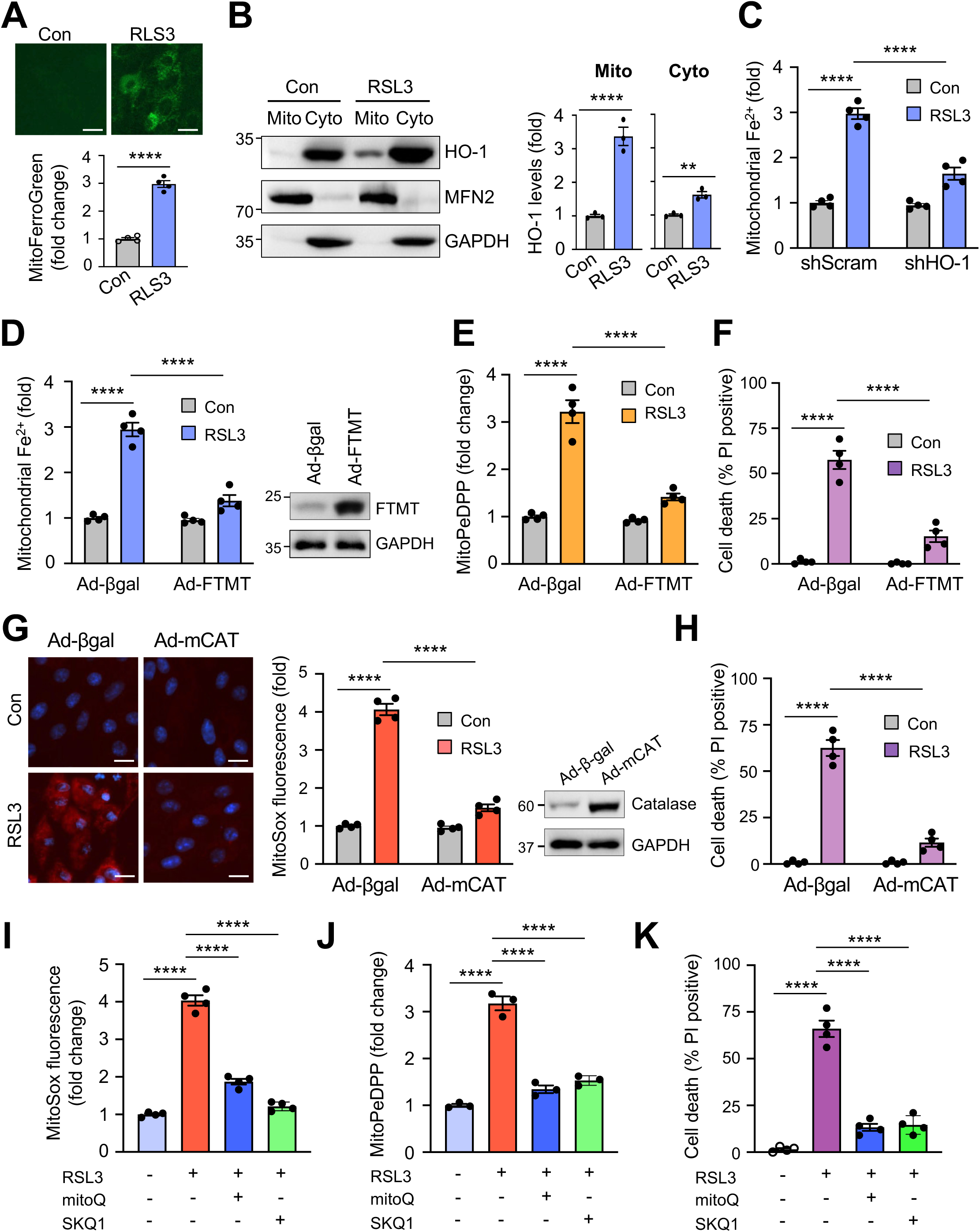
Inactivation of GPX4 promotes mitochondrial iron overload via HO-1 mitochondrial translocation. (**A**) Mitochondrial Fe^2+^ levels assessed with mito-FerroGreen in neonatal rat cardiomyocytes treated with 0.5 μM RSL3 or vehicle control for 8 h. n = 4. Scale bar, 25 μm. (**B**) Western blotting for the indicated proteins from mitochondrial and cytosolic fractions of cardiomyocytes treated with 0.5 μM RSL3 or vehicle control. n = 3. (**C**) Mitochondrial Fe^2+^ levels in cardiomyocytes infected with Ad-shHO-1 or Ad-shScram for 24 h followed by 0.5 μM RSL3 for 8 h. n = 4. (**D**) Mitochondrial Fe^2+^ levels in cardiomyocytes infected with adenoviral vectors encoding mitochondrial ferritin (Ad-FTMT) or Ad-βgal for 24 h followed by 0.5 μM RSL3 for 8 h. FTMT expression was assessed by Western blotting. n = 4. (**E**) Mitochondrial lipid peroxidation assessed with MitoPeDPP in cardiomyocytes treated as in D. (**F**) Quantification of cell death from cardiomyocytes infected with Ad-FTMT or Ad-βgal for 24 h followed by 0.5 μM RSL3 for 12 h. n = 4. (**G**) MitoSox fluorescence staining in cardiomyocytes infected with adenoviral vectors for mitochondria-targeted catalase (mCAT) or Ad-βgal for 24 h followed by 0.5 μM RSL3 for 8 h. n = 4. Scale bar, 25 μm. Catalase expression was assessed by Western blotting. (**H**) Quantification of cell death from cardiomyocytes infected with Ad-βgal or Ad-mCAT for 24 h followed by 0.5 μM RSL3 for 12 h. n = 4. (**I**-**K**) Measurements of MitoSox levels, mitoPeDPP, and cell death in cardiomyocytes pretreated with mitoQ, SKQ1, or vehicle control for 30 min followed by 0.5 μM RSL3 for 8 (I, J) or 12 h (K). n = 4. *\*\*P ≤* 0.01; *\*\*\*\*P ≤* 0.0001. Statistical analysis was performed using Student’s t-test (A, B), one-way ANOVA with Tukey’s post hoc test (I-K), or 2-way ANOVA with Tukey’s post hoc test (C-H).

### Targeted inhibition of mitochondrial iron overload attenuates cardiac ferroptosis and remodeling associated with *Gpx4* deficiency

To further determine the role of mitochondria in mediating cardiac ferroptosis and remodeling in vivo, we examined whether targeted inhibition of mitochondrial iron overload or ROS production prevents cardiac remodeling and dysfunction in *Gpx4* deficient mice. First, we assessed the effect of N,N′-bis (2-hydroxybenzyl) ethylenediamine-N,N′-diacetic acid (HBED), a lipophilic mitochondrial iron chelator^25^, and mitoTEMPO, a mitochondria-targeted antioxidant, on RSL3-induced ferroptosis in cardiomyocytes in vitro. Both HBED and mitoTEMPO effectively inhibited RSL3-induced lipid peroxidation and cell death in cardiomyocytes (**Fig. S7**). To further test this in vivo, *Gpx4*^fl/fl^MCM (KO) and MCM control mice were treated with HBED, mitoTEMPO, or vehicle control along with tamoxifen, followed by echocardiography and histological analysis (**Fig. 7**). Administration of both HBED and mitoTEMPO greatly improved cardiac dysfunction, ventricular dilation, and hypertrophy in *Gpx4*-deficient mice (**Fig. 7, A** to **E**). *ptgs2* mRNA and MDA levels were also normalized by HBED and mitoTEMPO (**Fig. 7, F** and **G**). These data further suggest that mitochondrial iron overload and lipid peroxidation critically mediate cardiac ferroptosis and remodeling induced by *Gpx4* deficiency.

**Figure 7.**
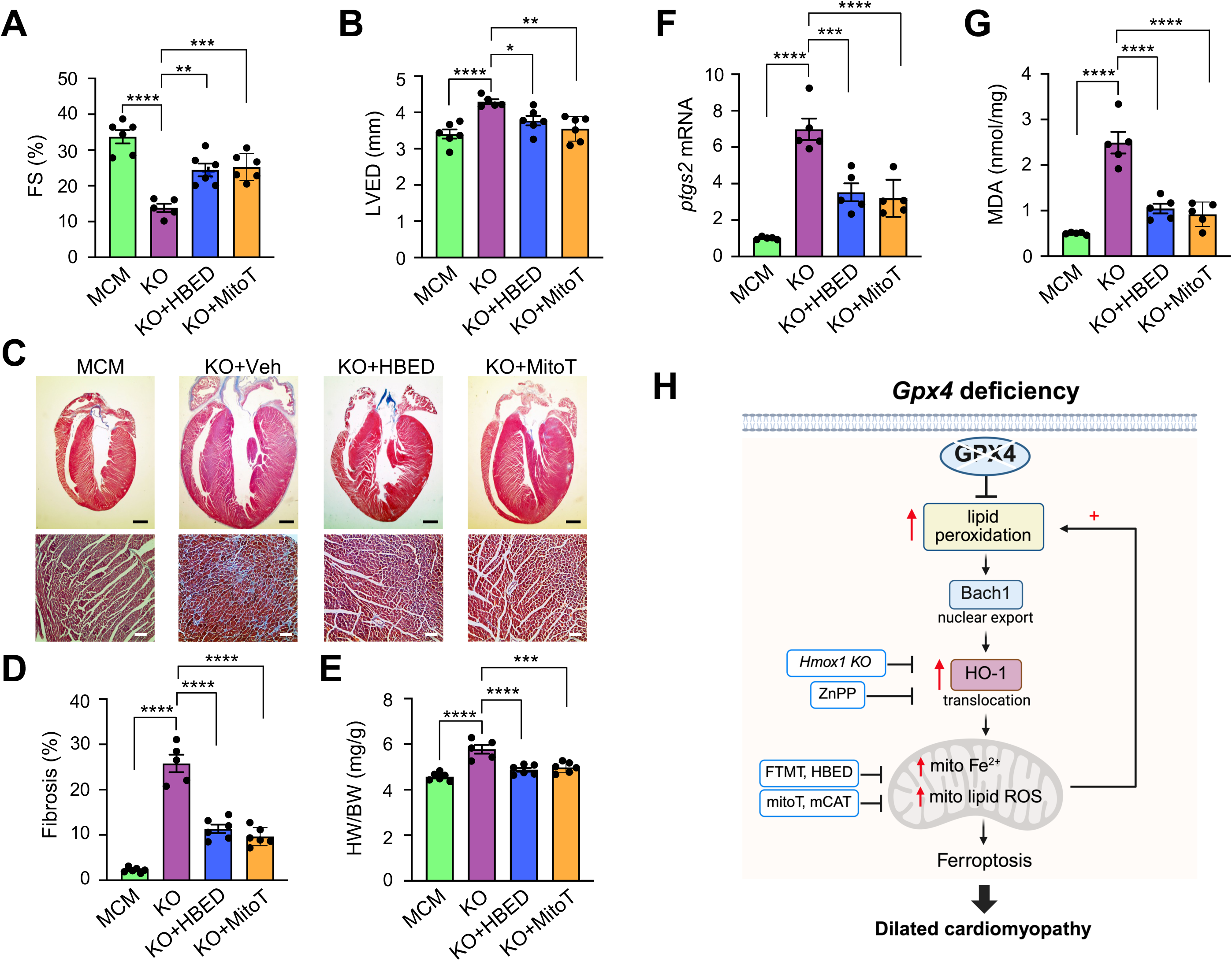
Targeted inhibition of mitochondrial iron overload or ROS accumulation attenuates cardiac remodeling and dysfunction in *Gpx4* deficient mice. (**A** and **B**) Fractional shortening (FS) and left ventricular end-diastolic dimension (LVED) measured by echocardiography from MCM and KO mice administrated with tamoxifen along with HBED (1 mg/kg/day, i.p.) or mitoTEMPO (5 mg/kg/day, i.p.) for 2 weeks. n = 5-7. (**C**) Masson’s trichrome-stained cardiac sections from mice treated as above. Scale bars: 1 mm in top panels; 50 μm in bottom panels. (**D**) Cardiac fibrosis in mice of the indicated genotypes. n = 5-7. (**E**) Heart weight to body weight ratio (HW/BW) from mice of the indicated genotypes. n = 5-7. (**F**) *ptgs2* mRNA levels in the heart of the indicated groups. n = 5-7. (**G**) MDA levels in cardiac extracts of the indicated groups. (**H**) Proposed model: Enhanced lipid peroxidation promotes mitochondrial iron overload and ferroptotic cell death via Bach1/HO-1 signaling. n = 5-7. *\*P ≤* 0.05; *\*\*P ≤* 0.01; *\*\*\*P ≤* 0.001; *\*\*\*\*P ≤* 0.0001. Statistical analysis was performed using one-way ANOVA with Tukey’s post hoc test.

## Discussion

Our results reveal a mechanistic link between lipid peroxidation and iron overload in ferroptosis via a Bach1-HO-1 dependent mechanism (**Fig. 7H**). Specifically, enhanced lipid peroxidation due to *Gpx4* deficiency promotes the nuclear export and degradation of Bach1, leading to upregulated HO-1 expression and increased iron release. HO-1 mediated iron overload further promotes lipid peroxidation via the Fenton reaction. Mitochondrial translocation of HO-1 serves as a key mechanism driving mitochondrial iron overload and ROS accumulation, which constitutes a positive feedback loop promoting lipid peroxidation and ferroptosis. Genetic and pharmacologic inactivation of HO-1 largely rescues the pathological phenotype of the *Gpx4* deficient heart by suppressing cellular iron overload. Moreover, targeted inhibition of mitochondrial iron overload or lipid peroxidation abrogates cardiac ferroptosis and dilated cardiomyopathy associated with *Gpx4* deficiency (**Fig. 7H**). These results indicate that enhanced lipid peroxidation due to *Gpx4* deficiency was sufficient to promote iron overload and ferroptosis, leading to pathological cardiac remodeling and dysfunction. Our data also identify Bach1-HO-1 signaling and mitochondrial iron overload as key mediators of ferroptosis in cardiomyocytes, which may represent new molecular targets for cardiac remodeling and heart failure.

The connection between lipid peroxidation and iron overload, two central events in ferroptosis, has not been directly investigated in previous studies. Here we found that enhanced lipid peroxidation intrinsically induces iron overload via an HO-1 dependent mechanism. HO-1 mediated iron overload has also been implicated in beta-thalassemia, sickle cell disease, and anthracycline cardiotoxicity.^26–28^ However, the role of HO-1 in ferroptosis remains controversial as both pro- and anti-ferroptotic roles of HO-1 have been reported, depending on cell types and ferroptotic inducers^20^. Similarly, both protective and detrimental effects of HO-1 are also observed in different disease settings.^14,28,29^ Impotently, we observed that ablation of *Hmxo-1* using a *Hmox1*^fl/+^αMHC-Cre mouse model elicited cardioprotection against adverse cardiac remodeling and heart failure following pressure overload (*Chen et al., unpublished observations*). Here, we further identified a dose-dependent, biphasic effect of HO-1 in regulating ferroptosis in cardiomyocytes. Our data support a model whereby moderate activation of HO-1 elicits a cytoprotective effect, whereas excessive and/or prolonged activation of HO-1 induces iron overload and ferroptosis. Of note, additional mechanisms may also contribute to iron overload during ferroptosis, such as NCOA4-mediated ferritinophagy, which promotes iron overload via ferritin iron release.^30^ However, we found that ablation of NCOA4 failed to inhibit iron overload or ferroptosis in cardiomyocytes following GPX4 inhibition. In line with this, a similar observation has also been reported in Hela cells^31^, suggesting that NCOA4-mediated ferritinophagy might be dispensable in this process.

Mechanistically, lipid peroxidation induces HO-1 expression in cardiomyocytes by promoting the nuclear export of Bach1, a transcriptional repressor.^32^ Although HO-1 is a NRF2 target gene^33^, inactivation of Bach1 has been shown to be a prerequisite for HO-1 induction, which is dominant over NRF2-mediated HO-1 transcription.^34^ Moreover, Bach1 inactivation can induce HO-1 expression without NRF2 nuclear accumulation.^34^ Consistent with this notion, we found that deletion of Bach1 was sufficient to induce HO-1 expression in cardiomyocytes, which was associated with enhanced ferroptosis. Conversely, overexpression of Bach1 inhibited RSL3-induced ferroptosis in cardiomyocytes. In contrast, overexpression of NRF2 inhibited, whereas deletion of NRF2 promoted, RSL3-induced ferroptosis. Moreover, the expression of NQO1, another target gene of NRF2, was unaffected by GPX4 inactivation. Therefore, these data suggest that GPX4 regulates HO-1 primarily through a Bach1-dependent mechanism. Together, these results identified Bach1-HO-1 signaling as a key driver of iron overload and ferroptosis.

Importantly, we further identified HO-1 mitochondria translocation as a previously undescribed mechanism driving mitochondrial iron overload during ferroptosis. Inactivation of HO-1 or targeted inhibition of mitochondrial iron overload effectively suppressed ferroptosis induced by GPX4 inactivation. Of note, HO-1 mitochondrial translocation has also been observed in cells undergoing hypoxia, leading to mitochondrial ROS accumulation and dysfunction.^35^ The precise mitochondria targeting sequence of HO-1 has yet to be identified, but it is speculated that a cryptic mitochondria-targeting signal may exit which is regulated by redox states.^35^ Therefore, the molecular mechanism underlying HO-1 mitochondrial translocation warrants further investigation. Moreover, additional mechanisms may contribute to mitochondria iron overload during ferroptosis. For example, mitochondrial iron uptake through iron transporters, such as mitoferrin-2 and the mitochondrial Ca^2+^ and Fe^2+^ uniporter (MCU), may also mediate mitochondrial iron overload.^36,37^ Nonetheless, our results highlight an indispensable role of mitochondrial iron overload in mediating ferroptosis.

Our results also highlight a key role for mitochondrial iron overload in the execution of ferroptosis. Indeed, mitochondrial iron overload and lipid peroxidation were robustly induced by GPX4 inactivation. Moreover, blockade of mitochondrial iron overload by FTMT overexpression markedly inhibited RSL3-induced mitochondrial lipid peroxidation and ferroptosis. Inhibition of mitochondrial iron overload with HBED, a lipophilic mitochondrial iron chelator, also largely rescued cardiac remodeling and dysfunction in *Gpx4*-deficient mice in vivo. Compared to other common iron chelators, HBED can cross mitochondrial membranes with a higher affinity to ferrous iron, a longer half-life, and a relatively safer toxicity index..^25,38,39^ Of note, HBED has passed phase I clinical trial and is shown to be safe to administer to humans.^40^ Targeted inhibition of mitochondrial ROS with mitoQ, SKQ1, or mCAT also inhibited RSL3-induced ferroptosis in cardiomyocytes. Moreover, the mitochondria-targeted antioxidant mitoTEMPO markedly ameliorated myocardial ferroptosis and remodeling in *Gpx4*-deficient mice in vivo. These results support a key role of mitochondrial iron overload and lipid ROS accumulation in mediating ferroptosis in cardiomyocytes. Given adverse effects associated with classic ferroptosis inhibitors^41,42^, targeting mitochondrial iron overload and/or lipid peroxidation may serve as a new anti-ferroptotic strategy for cardiac remodeling and heart failure.

## Materials and Methods

### Reagents

Ferrostatin-1, (1S,3R)-RSL3, and Evan’s blue dye were from Sigma-Aldrich. Liproxstatin-1, deferoxamine, zVAD-fmk, mitoTEMPO, mitoquinone mesylate (mitoQ), SKQ1, and ZnPP were from Cayman Chemical. MitoPeDPP and mito-FerroGreen were from Dojindo. Necrostatin-1s was from Cell Signaling Biotechnology. HBED was from Santa Cruz Biotechnology. MitoSOX, propidium iodide, and Hoechst 33342 were from Invitrogen. The following antibodies were used: anti-GPX4 (59735), anti-HO-1 (82206), anti-mitofusion-2 (9482), and anti-catalase (14097) from Cell Signaling Biotechnology; anti-Bach1 (sc-271211), and anti-GAPDH (sc-32233) from Santa Cruz Biotechnology; Anti-FTMT (PAD252Mu01) from Cloud-Clone Corp.; anti-GPX4 (MAB5457) from R&D Systems.

### Animal models

All experiments involving animals were approved by the Institutional Animal Care and Use Committees of the University of Washington, and all studies were carried out in accordance with the approved protocols. *Gpx4*-floxed (*Gpx4*^fl/fl^) mice were obtained from Dr. Qitao Ran at University of Texas Health Science Center at San Antonio.^10^ *Gpx4^fl/fl^* mice were crossed with αMHC-MerCreMer mice^18^ to generate cardiomyocyte-specific *Gpx4* knockout mice, which ablate all GPX4 isoforms (cytosolic, mitochondrial, and nuclear). *Hmox1*-floxed (*Hmox1^fl/fl^*) mice were kindly provide by Dr. Claude Piantadosi and Dr. Hagir Suliman at Duke University^23^, which were crossed with *Gpx4*^fl/fl^-αMHC-MerCreMer mice. Mice of both sexes aging from 2-3 months were used unless otherwise stated. No animals were excluded during experiments or data analysis. No randomization was used to allocate experimental units. Investigators were blinded to genotype/group allocation during the experiments. In some experiments, mice were treated with tamoxifen (50 mg per kg body weight, i.p. for 5 days or 0.4 g/kg tamoxifen in Teklad global 2016 base rodent diet, TD.130859 from Inotiv). Where indicated, mice were given a daily intraperitoneal injection of ferrostatin-1 (1 mg/kg), ZnPP (5 mg/kg), mitoTEMPO (1 mg/kg), HBED (20 mg/kg) or vehicle control.

### Echocardiography and transverse aortic constriction

Mice were anesthetized with 2% isoflurane by inhalation and scanning was performed with a VisualSonics Vevo 2100 imaging system as described previously.^43^ M-mode ventricular dimensions were averaged from 3-5 cycles. Fractional shortening (FS) was calculated using left ventricular systolic and diastolic dimensions (LVES and LVED, respectively): FS = [(LVED - LVES)/LVED] x 100 (%). Transverse aortic constriction (TAC) was performed to induce cardiac pressure overload in mice using a 27-gauge needle as previously described.^44^ Mice were anesthetized with inhaled 2% isoflurane and were intubated and respirated throughout. Sham-operated mice underwent the same procedure without aortic constriction. Doppler echocardiography was performed on mice after TAC to ensure equal pressure gradients across the aortic constriction.^43^ Pressure gradients (PGs; mm Hg) across the aortic constriction were calculated from the peak blood velocity (Vmax) (m/s) (PG = 4 × Vmax2).

### Masson’s trichrome staining, TUNEL, and serum cTnI measurement

For histological analysis, hearts were fixed in 10% formalin/phosphate-buffered saline and dehydrated for paraffin embedding. Fibrosis was detected with Masson’s trichrome staining on 5 μm paraffin sections. Blue collagen staining was quantified using Metamorph 6.1 software.^45^ Terminal deoxynucleotidyl transferase dUTP nick end labeling (TUNEL) in paraffin sections was performed with an ApopTag peroxidase in situ apoptosis detection kit (Millipore) according to the manufacturer’s instructions.^45^ Mouse cTnI serum levels were measured using an ELISA kit from Life Diagnostics, Inc. (West Chester, PA). Absorbance at 450 nm (sample) and 630 nm (reference) was measured with a BioTek Synergy 2 microplate reader (BioTek).

### Evans blue dye uptake assay

Mice were injected with Evan’s blue dye (EBD, 100 mg/ kg i.p.). and sacrificed 24 h later. The hearts were embedded in Optimal Cutting Temperature Compound (Tissue-Tek) and frozen in liquid nitrogen. Cardiac cryosections (5 μm) were fixed with 4% paraformaldehyde, washed in PBS, and stained with wheat germ agglutinin conjugated to FITC (WGA, Sigma-Aldrich #L4895) for 1 h at room temperature to visualize the plasma membrane. Images were captured with an EVOS FL digital fluorescence microscope to determine EBD-positive area as red auto-fluorescence.

### Quantitative PCR

Total RNA was isolated from the heart using RNeasy fibrous tissue mini kit (Qiagen). The mRNA levels were determined by quantitative reverse transcription PCR using SuperScript IV reverse transcriptase (Thermo Fisher Scientific) for reverse transcription and a PowerUp SYBR Green PCR master mix (Thermo Fisher Scientific) for the quantitative PCR reaction. The following PCR primers were used: *Ptgs2* forward 5′-CAGACAACATAAACTGCGCCTT-3′ and reverse 5′-GATACACCTCTCCACCAATGACC-3’; atrial natriuretic peptide (*Nppa*) forward 5′-GCCCTGAGTGAGCAGACTG and reverse 5′-GGAAGCTGTTGCAGCCTA-3’; brain natriuretic peptide (*Nppb*) forward 5′-GGACCAAGGCCTCACAAAAG-3′ and reverse 5’-AAAGAGACCCAGGCAGAGTC-3’; skeletal muscle α-actin (*Acta1*) forward 5’-AATGAGCGTTTCCGTTGC-3’ and reverse 5’-ATCCCCGCAGACTCCATAC-3’; and GAPDH forward 5′-ACTGAGCAAGAGAGGCCCTA and reverse 5′-TATGGGGGTCTGGGATGGAA. All data were normalized to the GAPDH mRNA content and are expressed as the fold increase over the control group.

### Cell culture

Neonatal rat cardiomyocytes were prepared from hearts of 1- to 2-day-old Sprague-Dawley rats as we previously described.^45^ To isolate cardiomyocytes, the ventricles were minced in HBSS prior to enzymatic digestion and then subjected to 5 rounds of enzymatic digestion using 0.05% pancreatin (Sigma-Aldrich) and 84 U/mL collagenase (Worthington). Cells were collected by centrifugation at 500 g for 5 minutes and resuspended in M199 medium. After separation from fibroblasts, enriched cardiomyocytes were plated on 1% gelatin-coated plates. Cells were grown in M199 medium supplemented with 2% bovine growth serum (Thermo Fisher Scientific, SH3054103), 100 U/mL penicillin-streptomycin, and 2 mM L-glutamine.

### Adenoviral vectors

Adenoviral vectors encoding GPX4 shRNA, HO-1 shRNA, Bach1 shRNA, FTMT shRNA, or a scrambled shRNA were generated using the BLOCK-iT Adenoviral RNAi Expression System (Invitrogen). The core target sequence for GPX4 shRNA: 5’-GCCAGGAAGTAATCAAGAAAT-3’; HO-1 shRNA: 5’-GCTGACAGAGGAACACAAAGA-3’; Bach1 shRNA: 5’-GCGTACACAATATCGAGGAAT-3’; NCOA4 shRNA: 5’-AGAAGGAAGCGATGCATAAAT-3’; FTMT shRNA: 5’-GCTTTACGCATCCTACGTGTA-3’; scrambled shRNA: 5′-GCCTTAGGTTGGTCGAGAAA-3′. To generate HO-1, Bach1, or FTMT adenoviral vectors, HO-1 (Addgene #74672), Flag-Bach1 (Addgene #159130) and FTMT-Flag (GenScript #OHu55907) were cloned into pENTR3C and pAd/CMV/V5-DEST using the ViraPower Adenoviral Expression System (Invitrogen). Ad-mCAT was from University of Iowa Vector Core. Ad-NRF2 and Ad-shNRF2 were described previously.^46^ Adenoviral infections were performed as described previously at a multiplicity of infection of 10 to 50 plaque forming units per ml or as indicated.

### Cell death assays

Cell death was measured using a Cell Meter Apoptotic and Necrotic Detection kit (ATT Bioquest,) as we previously described .^47^ Briefly, cells were incubated at 37 °C for 30 min with Apopxin Green for detection of phosphatidylserine on cell surface, propidium iodide or 7-ADD for labeling the nucleus of cells with membrane rupture, and CytoCalcein for labeling live cell cytoplasm. Cell death was analyzed with an EVOS FL digital fluorescence microscope (AMG) and a Muse cell analyzer (MilliporeSigma). Cell viability was assessed using the Muse Count & Viability assay kit and a Muse cell analyzer (MilliporeSigma). All imaging data are representative of at least three randomly selected fields.

### Western blotting

Cardiac tissue or cultured cells were lysed in RIPA buffer (20 mM Tris pH 7.5, 200 mM NaCl, 1% Triton-X 100, 2 mM DTT, 5 mM p-nitrophenyl phosphate, 2 mM sodium orthovanadate, 1X protease inhibitor cocktail (Roche). Equal amounts of protein were subjected to SDS-PAGE and transferred to PVDF membranes (EMD Millipore, IPFL00010). Western blotting followed by enhanced chemiluminescence detection was performed as previously described.^47^

### Malondialdehyde (MDA) measurement

MDA was measured using the thiobarbituric acid reactive substances (TBARS) assay kit (Cayman Chemical). Briefly, thiobarbituric acid (TBA) was reacted with MDA in the samples at 100°C for 60 min, and MDA-TBA adduct was fluorometrically measured using a BioTek Synergy 2 microplate reader at an excitation wavelength of 530 nm and an emission wavelength of 550 nm.

### Cardiac total tissue iron, non-heme iron, and heme levels

Total tissue iron in the heart was measured with a ferrozine-based colorimetric assay kit (AdipoGen). Cardiac non-heme iron was measured using a bathophenanthroline-based colorimetric assay as previously described.^48^ Cardiac heme levels were measured using a QuantiChrom heme assay kit (BioAssay Systems).

### Labile iron measurement

The levels of labile iron were measured by the calcein-AM method as previously described.^49^ Briefly, cells were incubated with 1 μM calcein-AM at 37°C for 10 min followed by three washes with PBS. The fluorescence was measured using a BioTek Synergy 2 fluorescence microplate reader. Cells were then treated with 100 μM 2’,2’-bipyridine (BIP) at 37°C for 10 min and the fluorescence was measured again. The changes in fluorescence upon BIP treatment was used to determine the levels of labile iron.

### Cytosolic and mitochondrial fractions

Cytosolic and mitochondrial fractions were prepared as previously described.^50^ Briefly, cells were suspended in sucrose-mannitol buffer (20 mM HEPES, pH 7.5, 2 mM EDTA, 70 mM sucrose, 220 mM mannitol, 5 mM NaF, protease inhibitor cocktail (Roche) and homogenized using a Teflon homogenizer. The homogenates were centrifuged at 600 g for 10 min at 4 °C. The supernatant was re-centrifuged at 10,000 g for 15 min at 4 °C to collect the supernatant (cytosolic fraction) and pellet (mitochondrion fraction). The purity of cytosolic and mitochondrial fractions was validated by Western blotting using anti-GAPDH and anti-mitofusin-2 antibodies, respectively.

### Mitochondrial ferrous iron

Mitochondrial iron was measured using Mito-FerroGreen (Dojindo), a fluorescence probe for ferrous ion (Fe^2+^) in the mitochondria.^13^ Cells were incubated in 5 μM Mito-FerroGreen for 30 minutes at 37°C. After three washes in PBS, ammonium iron sulfate (100 μM) was added to the cells. Mitochondrial iron was fluorometrically measured at an excitation wavelength of 505 nm and an emission wavelength of 535 nm using a BioTek Synergy 2 microplate reader or visualized using an EVOS FL digital fluorescence microscope.

### Mitochondrial lipid peroxidation

Mitochondrial lipid peroxidation was assessed using MitoPeDPP (Dojindo), a fluorescence probe that specifically detects lipid peroxides in the mitochondrial inner membrane.^13^ Cells were incubated with 0.5 μM MitoPeDPP solution for 30 minutes at 37°C. Following three washes with PBS, mitochondrial lipid peroxidation was fluorometrically measured using a BioTek Synergy 2 microplate reader at an excitation wavelength of 452 nm and an emission wavelength of 470 nm.

### Mitochondrial ROS

MitoSOX Red (Invitrogen) was used for analyzing mitochondrial reactive oxygen species (ROS). Cells were loaded with MitoSOX at 5 μM for 30 min at 37 °C. After washing three times with PBS, fluorescence was detected by an EVOS FL digital fluorescence microscope (AMG) and quantified with ImageJ software.

### Statistical analysis

Results are presented as mean ± SEM. Sample number was predetermined based on our previous results from similar studies followed by power analysis (α = 0.05; power = 80%). Statistical analysis was performed using GraphPad Prism 9 (GraphPad, San Diego, CA). Student’s two-tailed t test was used for comparison between 2 groups. Comparisons between multiple groups were made using one-way or two-way analysis of variance (ANOVA) with Tukey’s post hoc test. Mann-Whitney U-test or Kruskal-Wallis test followed by post hoc Mann-Whitney U-test with Bonferroni’s correction was used for studies with small sample sizes. The log rank test was used for survival studies. *P* < 0.05 were considered significant. All surgeries, histological analysis, and echocardiographic measurements were performed blinded.

## Acknowledgements

*Gpx4*-floxed mice were kindly provided by Dr. Qitao Ran at University of Texas Health Science Center at San Antonio. We thank Dr. Claude Piantadosi and Dr. Hagir Suliman at Duke University for providing *Hmox1*-floxed mice. We also thank Dr. Jeffery Molkentin at the Cincinnati Children’s Hospital for providing αMHC-MerCreMer mice.

---

**Supplementary Figure 1.**
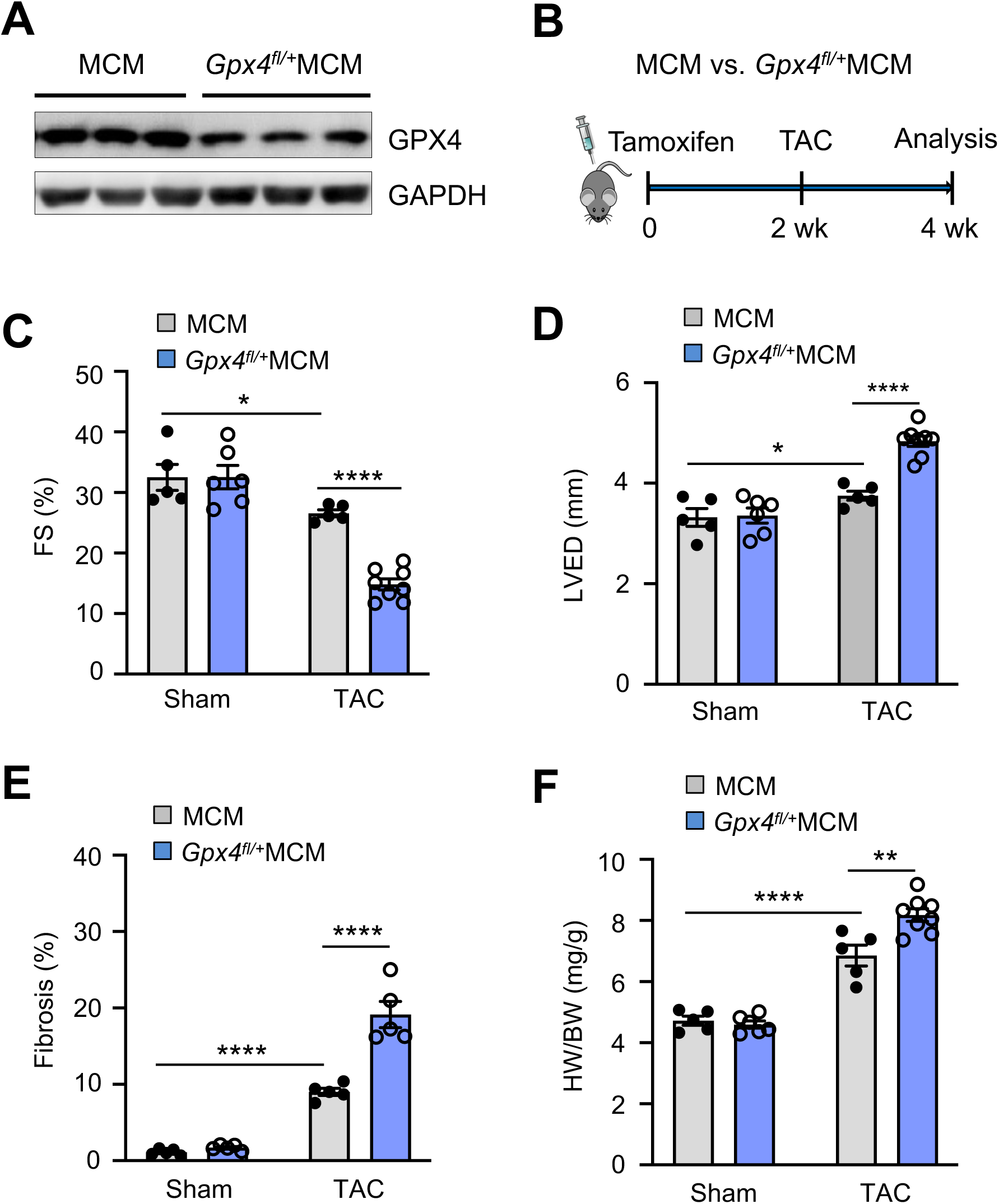
G*p*x4 deficient mice were predisposed to cardiac remodeling and dysfunction following pressure overload. (**A**) Western blotting for GPX4 and GAPDH in the mouse heart following tamoxifen administration (50 mg/kg/day for 5 days, i.p.). (**B**) Experimental regime of tamoxifen administration (50 mg/kg/day for 5 days, i.p.) followed by transverse aortic constriction (TAC) and echocardiographic/histological analysis. (**C** and **D**) Fractional shortening and left ventricular end diastolic dimension in MCM and *Gpx4*fl/+MCM mice after sham or TAC surgery. (**E**) Cardiac fibrosis. (**F)** Heart weight to body weight ratio. n = 5-8. \**P* ≤ 0.05; \*\**P* ≤ 0.01; \*\*\*\**P* ≤ 0.0001. Statistical analysis was performed using 2-way ANOVA with Tukey’s multiple-comparison test.

**Supplementary Figure 2.**
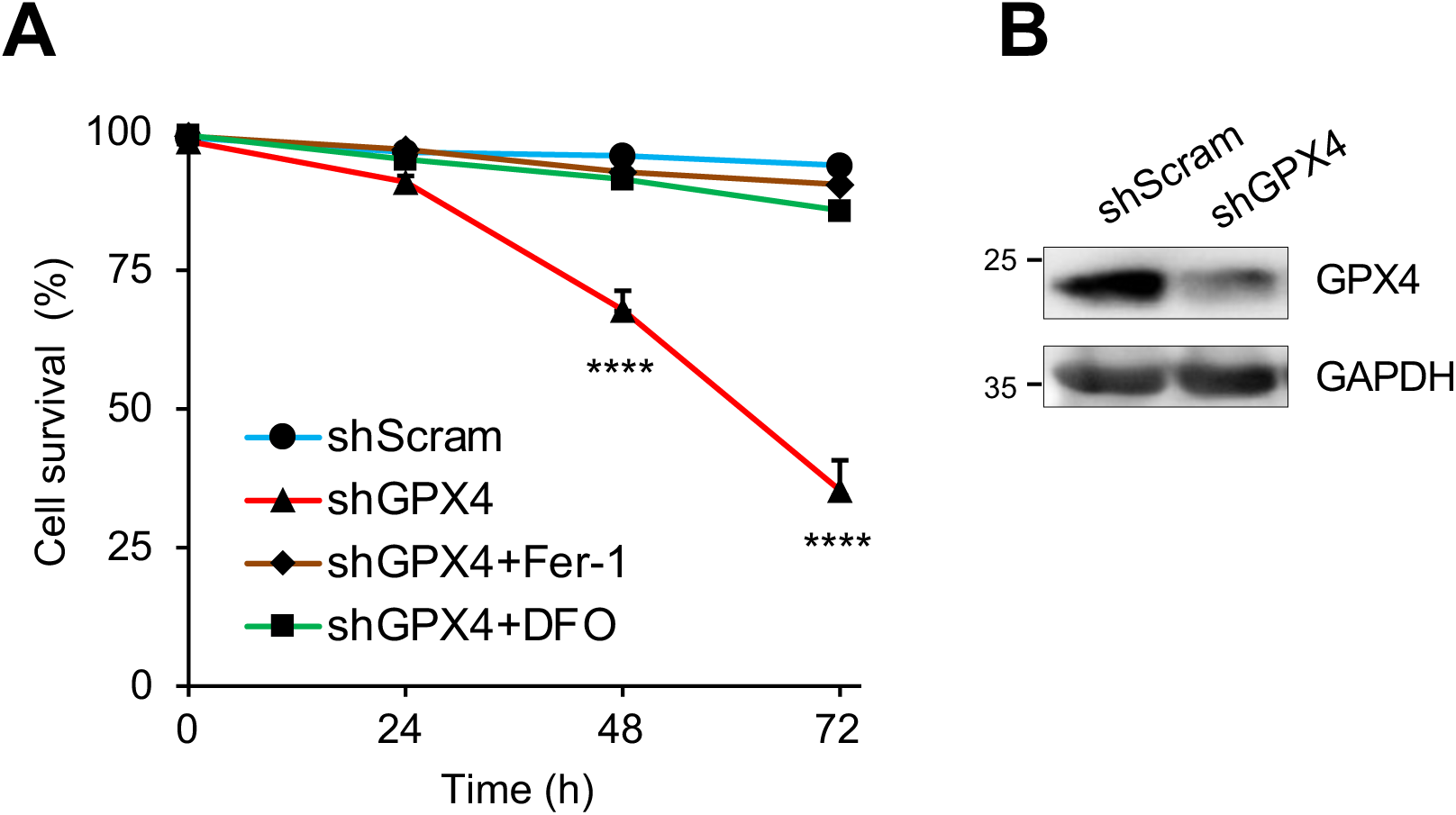
Loss of GPX4 induced ferroptosis in cardiomyocytes. (**A**) Cell survival rate in neonatal cardiomyocytes infected with adenoviral vectors encoding shGPX4 or a scrambled sequence (shScram) in the presence or absence of 10 μM Fer-1 (ferrostatin-1, an inhibitor of lipid peroxidation and ferroptosis) or 10 μM DFO (deferoxamine, an iron chelator) for 0 - 72 h. (**B**) Western blotting for the indicated proteins in cardiomyocytes subjected to adenoviral infection for 72 h. n = 3. \*\*\*\**P* ≤ 0.0001. Statistical analysis was performed using one-way ANOVA with Tukey’s post hoc test.

**Supplementary Figure 3.**
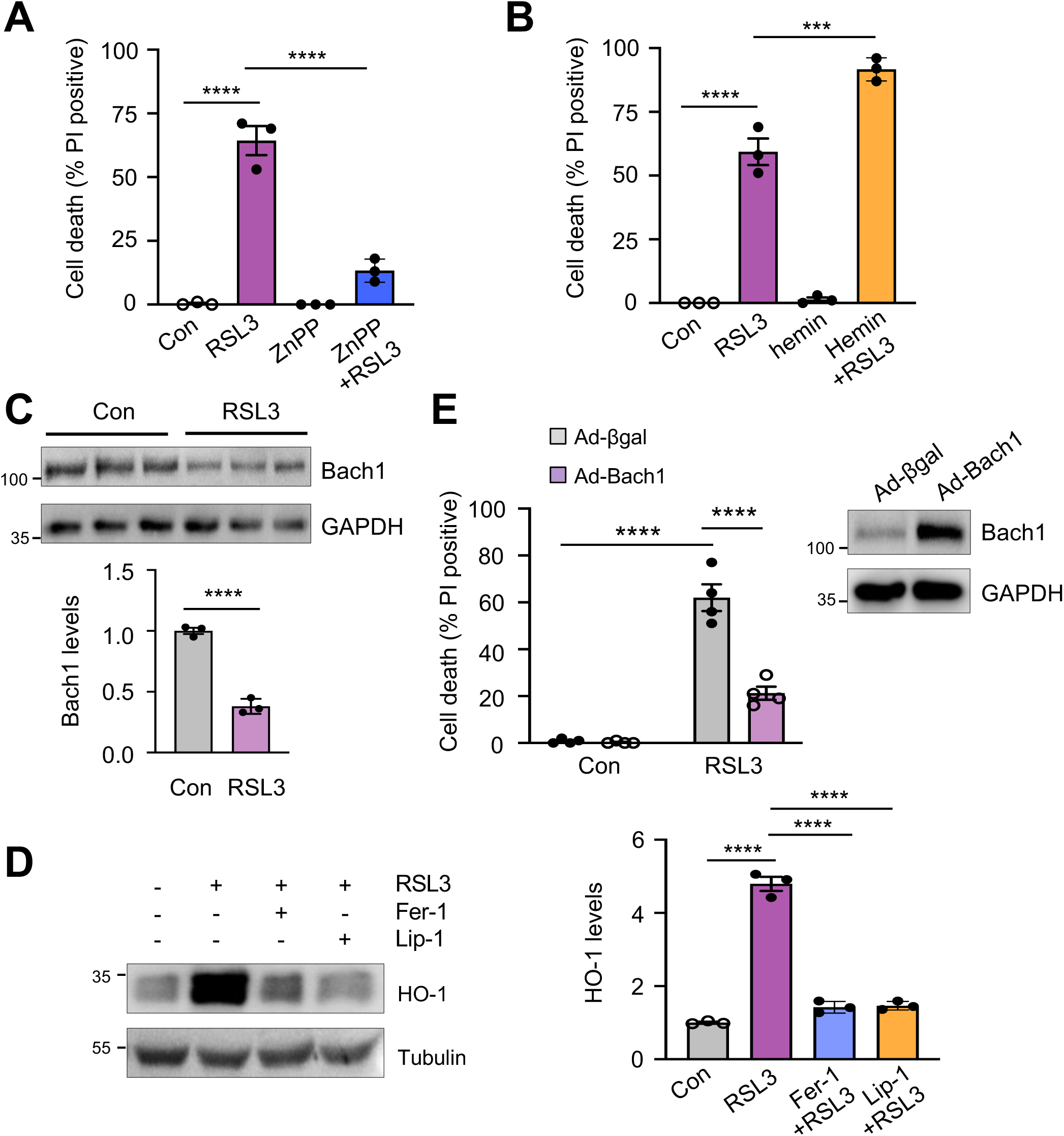
Bach1-HO-1 signaling regulates ferroptosis. (**A**) Cell death in neonatal cardiomyocytes treated with 0.5 μM RSL3 or vehicle control with or without 5 μM ZnPP for 12 h. n = 3. (**B**) Cell death in neonatal cardiomyocytes treated with RSL3 or vehicle control with or without 10 μM hemin for 12 h. n = 3. (**C**) Western blot and quantification of Bach1 in cardiomyocytes treated with RSL3 for 8 h. n = 3. (**D**) Western blot and quantification of HO-1 in neonatal cardiomyocytes treated with RSL3 or vehicle control in the presence of 10 μM Fer-1 or 1 μM Lip-1 for 8 h. n = 3. (**E**) Cell death in neonatal cardiomyocytes infected with the indicated adenoviral vectors for 24 h followed by RSL3 or vehicle control for 12 h. n = 4. \*\*\**P* ≤ 0.001; \*\*\*\**P* ≤ 0.0001.Statistical analysis was performed using Student’s t-test (**C**), one-way ANOVA with Tukey’s post hoc test (**A, B, D**), or 2-way ANOVA with Tukey’s post hoc test (**E**).

**Supplementary Figure 4.**
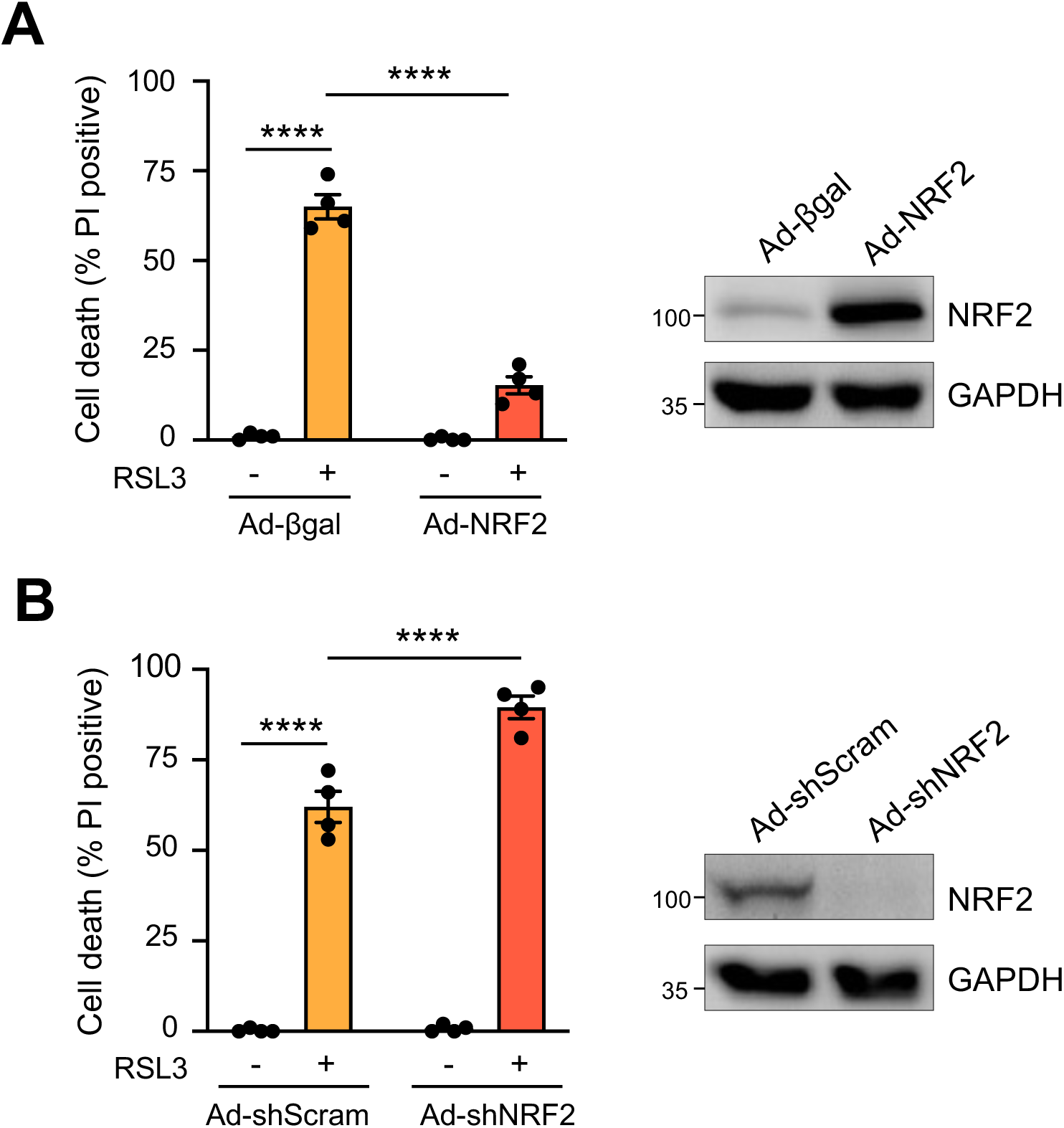
NRF2 negatively regulates RSL3-induced ferroptosis. (**A**) Cell death in neonatal cardiomyocytes infected with the indicated adenoviral vectors for 24 h followed by treatment with 0.5 μM RSL3 or vehicle control for 12 h. n = 4. (**B**) Cell death in neonatal cardiomyocytes infected with the indicated adenoviral vectors for 48 h followed by 0.5 μM RSL3 or vehicle control for 12 h. n = 4. \*\*\*\**P* ≤ 0.000. Statistical analysis was performed using 2-way ANOVA with Tukey’s post hoc test.

**Supplementary Figure 5.**
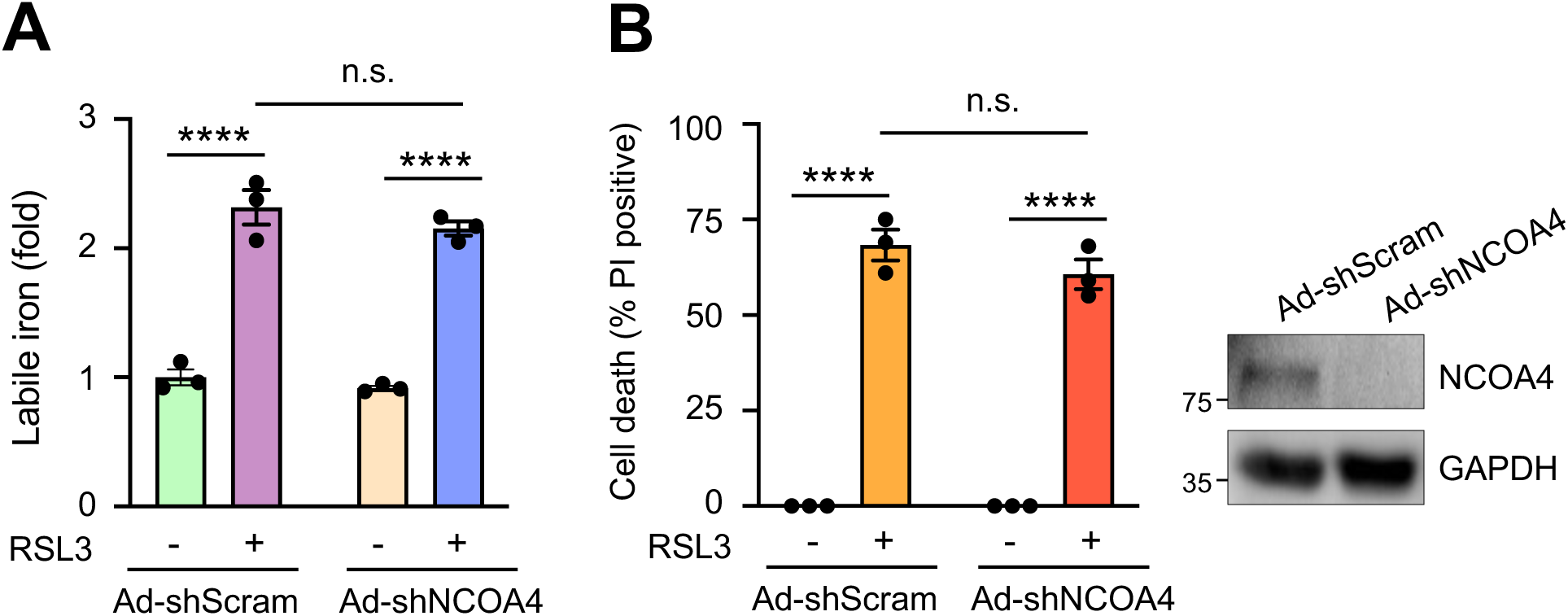
Deletion of NCOA4 had minimal effects on RSL3-induced iron accumulation and ferroptosis. (**A** and **B**) Labile iron levels and cell death were assessed in neonatal cardiomyocytes infected with the indicated adenoviral vectors for 48 h followed by treatment with 0.5 μM RSL3 or vehicle control for 12 h. n = 3. \*\*\*\**P* ≤ 0.0001. n.s., not significant. Statistical analysis was performed using 2-way ANOVA with Tukey’s post hoc test.

**Supplementary Figure 6.**
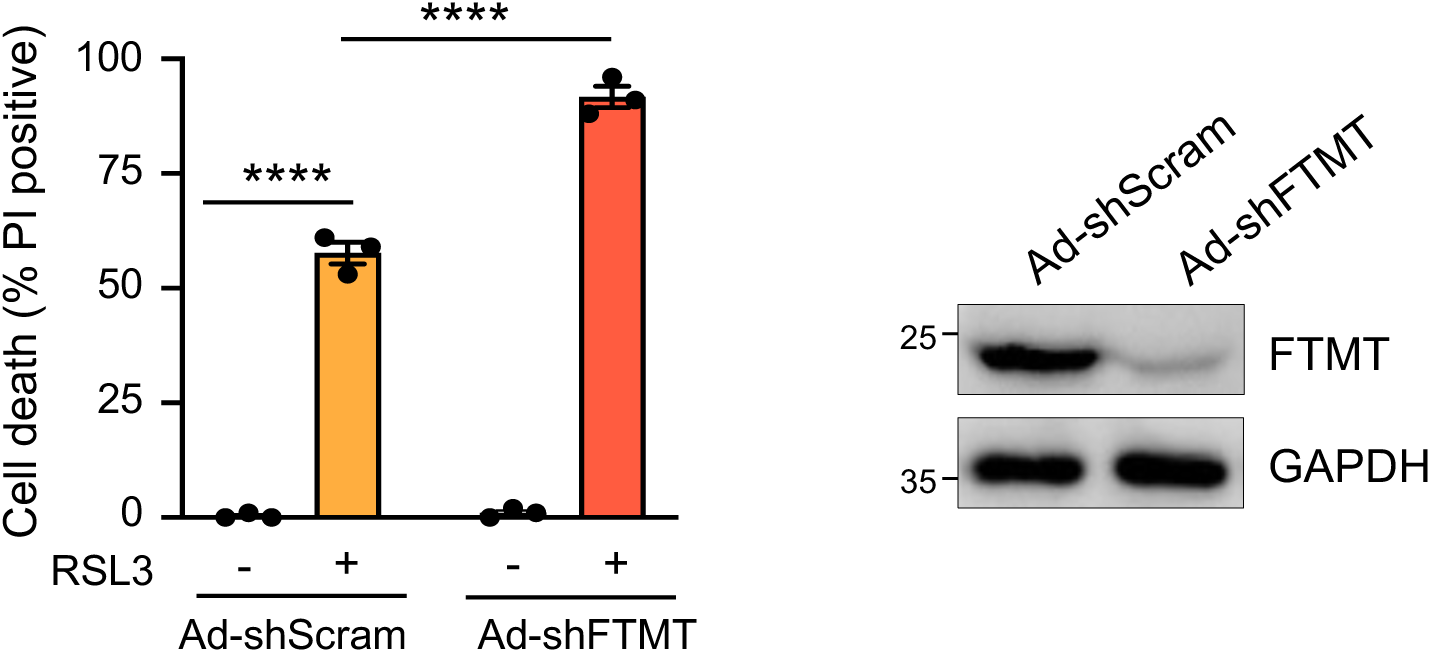
Deletion of FTMT promoted RSL3-induced ferroptosis. Cell death in neonatal cardiomyocytes infected with the indicated adenoviral vectors for 48 h followed by treatment with 0.5 μM RSL3 or vehicle control for 12 h. n = 3. \*\*\*\**P* ≤ 0.0001Statistical analysis was performed using 2-way ANOVA with Tukey’s post hoc test.

**Supplementary Figure 7.**
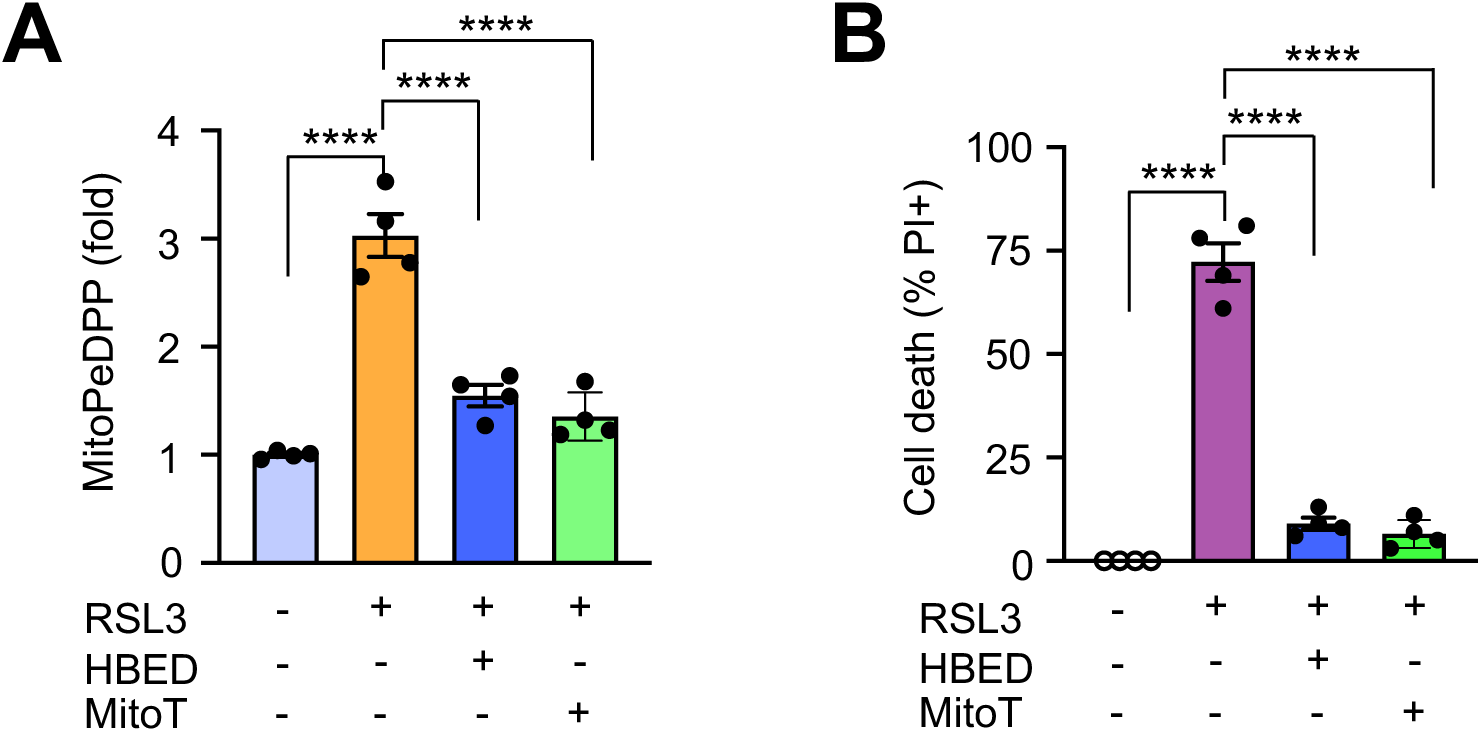
Blockade of mitochondrial iron overload or ROS accumulation prevented RSL3-induced ferroptosis. (**A** and **B**) Mitochondrial lipid peroxidation (mitoPeDPP) and cell death in neonatal cardiomyocytes pretreated with HBED, mitoTEMPO (mitoT), or vehicle control for 30 min followed by 0.5 μM RSL3 for 8 h (A) or 12 h (B). n = 4. \*\*\*\**P* ≤ 0.0001. Statistical analysis was performed using one-way ANOVA with Tukey’s post hoc test.

